# TBXT dose sensitivity and the decoupling of nascent mesoderm specification from EMT progression in 2D human gastruloids

**DOI:** 10.1101/2023.11.06.565933

**Authors:** Emily A. Bulger, Ivana Muncie-Vasic, Ashley R.G. Libby, Todd C. McDevitt, Benoit G. Bruneau

**Author notes:** Current address: Vitra Labs, San Francisco, CA. Current address: The Francis Crick Institute, London, NW1 1AT, United Kingdom (Ashley R.G. Libby). Current address: Genentech, South San Francisco, CA. Corresponding Author: Benoit G. Bruneau Keywords: mesoderm, gastrulation, brachyury, axial elongation, gastruloid, EMT.

## Abstract

In the nascent mesoderm, levels of Brachyury (TBXT) expression must be precisely regulated to ensure cells exit the primitive streak and pattern the anterior-posterior axis, but how this varying dosage informs morphogenesis is not well understood. In this study, we define the transcriptional consequences of TBXT dose reduction during early human gastrulation using human induced pluripotent stem cell (hiPSC)-based models of gastrulation and mesoderm differentiation. Multiomic single-nucleus RNA and single-nucleus ATAC sequencing of 2D gastruloids comprised of WT, TBXT heterozygous (TBXT-Het), or TBXT null (TBXT-KO) hiPSCs reveal that varying TBXT dosage does not compromise a cell’s ability to differentiate into nascent mesoderm, but that the loss of TBXT significantly delays the temporal progression of the epithelial to mesenchymal transition (EMT). This delay is dependent on TBXT dose, as cells heterozygous for TBXT proceed with EMT at an intermediate pace relative to WT or TBXT-KO. By differentiating iPSCs of the allelic series into nascent mesoderm in a monolayer format, we further illustrate that TBXT dose directly impacts the persistence of junctional proteins and cell-cell adhesions. These results demonstrate that EMT progression can be decoupled from the acquisition of mesodermal identity in the early gastrula and shed light on the mechanisms underlying human embryogenesis.

## INTRODUCTION

In the early embryo, cells must precisely regulate gene expression to ensure the organism progresses through standard hallmarks of development. One key developmental timepoint in vertebrate embryogenesis is the establishment and morphogenesis of the primitive streak (PS), a transient structure of the posterior embryo that initiates germ layer formation and establishes bilateral symmetry (Mikawa et al., 2004). Cells follow precisely orchestrated migration patterns as they ingress into the streak, undergo an epithelial-to-mesenchymal transition (EMT) to delaminate from the streak, and expand outward as individual mesenchymal cells to form endodermal and mesodermal germ lineages (Ciruna & Rossant, 2001; Dale et al., 2006; Schoenwolf & Smith, 2000). Timing of epithelial cell ingression into the PS informs cell fate, as cells ingressing early during gastrulation give rise to cranial mesoderm (Lawson et al., 1991), while subsequently ingressing cells form axial, paraxial, and lateral mesoderm of the anterior trunk (Wilson & Beddington, 1996). Posterior trunk mesoderm, including caudal somites, emerges later from a separate progenitor population in the tailbud known as the neuromesodermal progenitors (NMPs) (Henrique et al., 2015; Tzouanacou et al., 2009).

The transcription factor Brachyury (TBXT (human), also T/Bra (mouse)) has a conserved role in mesoderm differentiation across vertebrates (Technau, 2001) and is widely utilized as one of the first markers of nascent mesoderm. *T/Bra* is initially expressed in the posterior embryo just before the emergence of the PS, and as gastrulation progresses its expression domain expands to include the PS, notochord, and later the tailbud NMP population (RiveraLPérez & Magnuson, 2005; Wilkinson et al., 1990). Still, how precisely *T/Bra* orchestrates the interplay between germ layer specification and PS morphogenesis is unclear. Human stem-cell-based models show TBXT is required for mesoderm induction (Bernardo et al., 2011; Faial et al., 2015) and studies in Xenopus show that the TBXT homolog, XBra, can induce different mesodermal cell types in a dose-dependent manner (Faial et al., 2015; O’Reilly et al., 1995). In contrast, mice with loss-of-function *T/Bra* mutations appear to generate an early mesoderm population (Hashimoto et al., 1987; Rashbass et al., 1991; Wilson et al., 1995; Yanagisawa et al., 1981). Morphogenesis, however, is dramatically altered in these mutants, as *T/Bra^-/-^* mesoderm-like cells lose their ability to properly migrate away from the embryonic midline and accumulate in the node, PS, and regions immediately ventral to the streak. This aberrant cellular distribution, coupled with later-onset defects in tailbud mesoderm specification, ultimately causes mutant embryos to not develop a notochord, have significant body axis truncations rostral to somite seven, and die from incomplete allantois development. How *T/Bra* loss drives this phenotype, including if and how its mode of misregulation is conserved in humans, has not been fully defined.

Interestingly, this mutant axial truncation phenotype appears to be correlated to *T/Bra* dose. Mice heterozygous for *T/Bra* show a small but notable accumulation of cells in the same domains that see cell accumulation in *T/Bra* homozygous knock-out mice (Wilson & Beddington, 1997; Wilson et al., 1993, 1995). Heterozygous mice are viable but later generate short tails and can display notochord and sacral malformations (Chesley, 1935; DobrovolskaiaLZavadskaia, 1927; Stott et al., 1993). In humans, hypomorphic *TBXT* expression manifests as partial absences or abnormal fusions of the tailbone, pelvis, or lower vertebrae, and, like in mouse, *TBXT*-loss-of-function mutations are embryonic lethal (Chen et al., 2023; Ghebranious et al., 2008; Papapetrou et al., 1999; Postma et al., 2014). The intermediate axial truncation phenotype seen in both human and mouse when TBXT expression is reduced suggests that the mechanism governing the spatial patterning of mesoderm once it exits the vertebrate PS is dependent on precisely calibrated expression levels of *TBXT*.

For TBXT+ cells to become motile and exit the PS, they must undergo EMT, including restructuring cell-cell adhesions, the cytoskeleton, and the composition of the extracellular matrix (ECM). In early amniote gastrulation, EMT is tightly associated with the acquisition of mesoderm fate, and it has been suggested that EMT initiates mesoderm commitment in hESCs (Evseenko et al., 2010). TBXT also promotes EMT in the context of cancer (Fernando et al., 2010; Roselli et al., 2012). Still, how TBXT dosage modulates EMT in the context of mesoderm commitment to ensure cells acquire increased motility is not fully defined.

Acquisition of cell fate preceding and throughout the establishment of the primitive streak is primarily controlled by a network consisting of BMP, WNT, and Nodal signaling (Arnold & Robertson, 2009). This network can be manipulated *in vitro* to yield hESC colonies, termed 2D gastruloids, that reproducibly generate concentric rings of epiblast, mesoderm, endoderm, and extraembryonic cells from the center outwards (Minn et al., 2020, 2021; Warmflash et al., 2014). 2D gastruloids have been applied to understanding the minimal inputs driving multicellular patterning and migration kinetics throughout the course of gastrulation, including PS morphogenesis (Joy et al., 2021; Libby et al., 2019; Martyn et al., 2019).

Here, we adapt this 2D micropatterned gastruloid culture to investigate how TBXT dose-dependently controls mesodermal cell identity, EMT, and subsequent migratory behavior during early human gastrulation. We demonstrate that varying levels of TBXT expression modulate human PS morphogenesis by controlling the timing of EMT, including the persistence of junctional proteins and cell-cell adhesions, without compromising a cell’s ability to differentiate into nascent mesoderm. We conclude that initial mesoderm specification can be decoupled from the temporal progression of EMT during early gastrulation, thus significantly improving our understanding of cell fate acquisition and morphogenesis during early human development.

## RESULTS

### Generation of hiPSC TBXT allelic series and 2D Gastruloids

To investigate the effect of TBXT dose on PS morphogenesis, we engineered a WTC11-LMNB1-GFP-derived hiPSC allelic series: *TBXT*^+/+^ (WT), *TBXT*^+/-^ (TBXT-Het), and *TBXT*^-/-^ (TBXT-KO), by targeting the first exon of TBXT with CRISPR/Cas9 (Fig. 1A, S1). The resulting indel created a premature stop codon in one or two alleles, respectively. The WT line used in subsequent experiments was derived from a subclone that was exposed to the TBXT sgRNA but remained unedited. Conducting a western blot for TBXT in a monolayer of cells exposed to the WNT pathway agonist CHIR99021 for 48 hours revealed the expected decrease in TBXT expression across the allelic series (Figs. 1B-B’). Of note, the TBXT protein level in the TBXT-Het was approximately 80% of the WT protein level, suggesting intrinsic dose compensation mechanisms are active in the TBXT gene regulatory network (Fig. 1B’). Knock-out efficiency was further confirmed through immunofluorescence (IF), which demonstrated the complete absence of TBXT protein in the TBXT-KO (Figs. 1D).

**Figure 1:**
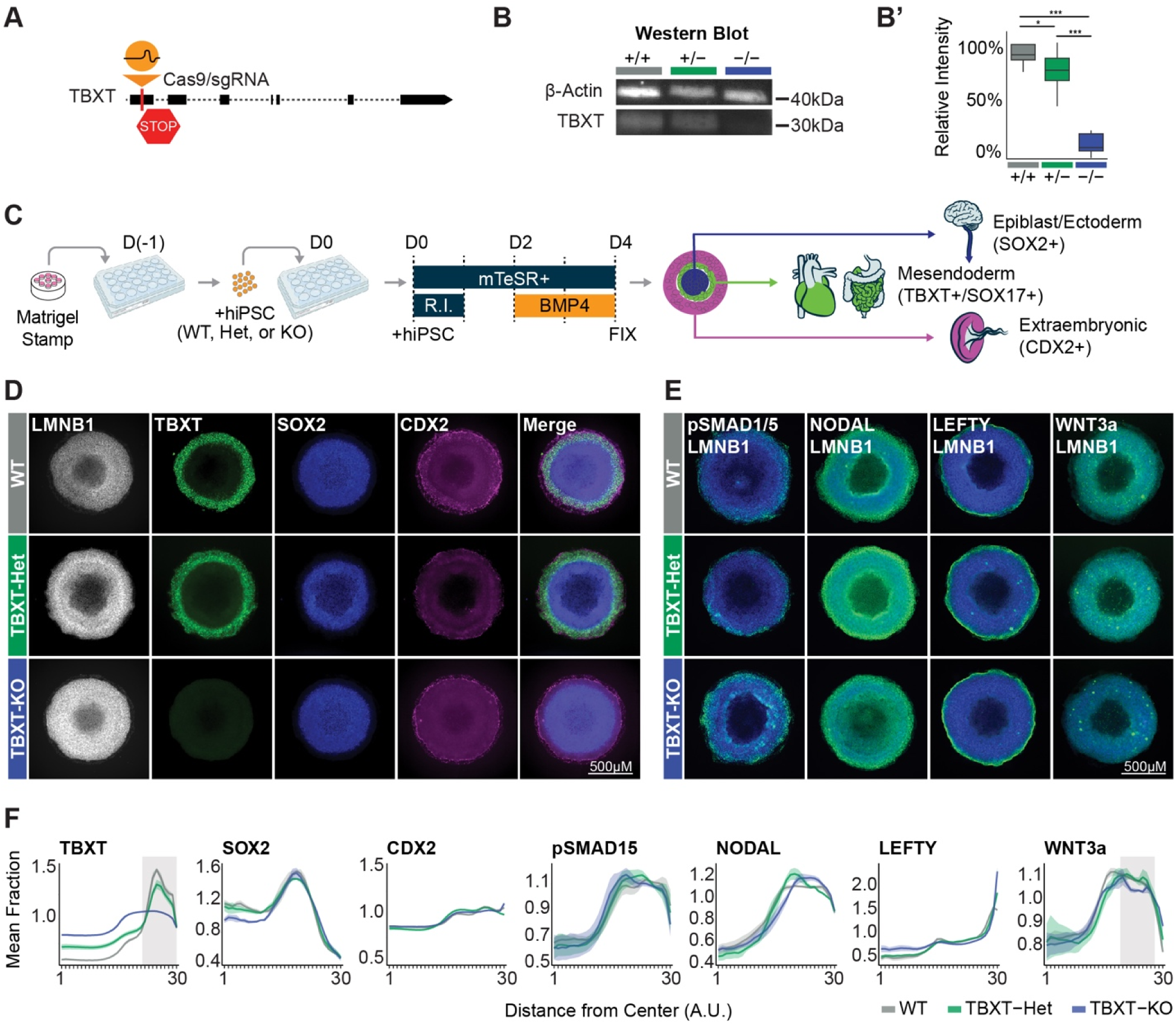
TBXT dose does not dramatically impact lineage emergence in 2D gastruloids. (**A**) Schematic of the location within the TBXT locus targeted by sgRNA to generate TBXT-Het and TBXT-KO. (**B**) Western blot and (**B’**) image intensity quantification showing the knock-out efficiency of TBXT sgRNA across the allelic series (*p<0.5, **p<0.1, ***p<0.01 by unpaired t-test, n = 5). (**C**) Schematic depicting the 2D gastruloid differentiation strategy. (**D-E**) Immunofluorescent images of gastruloids for canonical gastrulation markers. “R.I.” indicates ROCK inhibitor. (**E**) Green staining indicates protein of interest, while blue staining indicates LMNB1 (**F**) Quantification of fluorescent intensity of canonical gastrulation markers (n = TBXT: 5 WT, 15 TBXT-Het, 10 TBXT-KO; SOX2: 4 WT, 3 TBXT-Het, 3 TBXT-KO; CDX2: 5 WT, 15 TBXT-Het, 10 TBXT-KO; Lefty: 6 WT, 7 TBXT-Het, 8 TBXT-KO; Nodal: 8 WT, 3 TBXT-Het, 6 TBXT-KO; pSMAD1/5: 8 WT, 7 TBXT-Het, 4 TBXT-KO; WNT3A: 8 WT, 7 TBXT-Het, 4 TBXT-KO). Shadows around data indicate SEM. Gray vertical highlights indicate regions of interest based on variability in expression.

### TBXT dose does not dramatically impact lineage emergence in 2D gastruloids

To begin to dissect how TBXT dose shapes the earliest stages of nascent mesoderm morphogenesis, we subjected the allelic series to 2D gastruloid differentiation, which reproducibly generates concentric rings of radially patterned primary germ layers, primordial germ cell-like cells (PGCLCs), and extraembryonic-like cells after 48 hours of BMP4 exposure (Minn et al., 2020; Warmflash et al., 2014) (Fig.1C, Methods). We then conducted IF in colonies of each genotype to identify if and how TBXT expression broadly influences germ layers’ patterning, proportion, and identity during early human gastrulation.

Using antibodies targeting epiblast/ectoderm (SOX2+), mesoderm (EOMES+/SOX17-), endoderm/PGCLCs (SOX17+), and extraembryonic cells (CDX2+), we observed a conserved spatial distribution of the germ layers between WT, TBXT-Het, and TBXT-KO colonies (Figs. 1D, F, S2). The consistent expression pattern of these broad germ layer markers suggests that the intrinsic BMP ➔ WNT➔ NODAL feedback loop is sustained in the absence of TBXT as predicted *in silico* (Kaul et al., 2023), and reflects the ability of nascent mesoderm to develop in murine *T/Bra*^-/-^ models during the initial stages of gastrulation (Beddington et al., 1992; Rashbass et al., 1994; Wilson et al., 1995).

**Figure 2:**
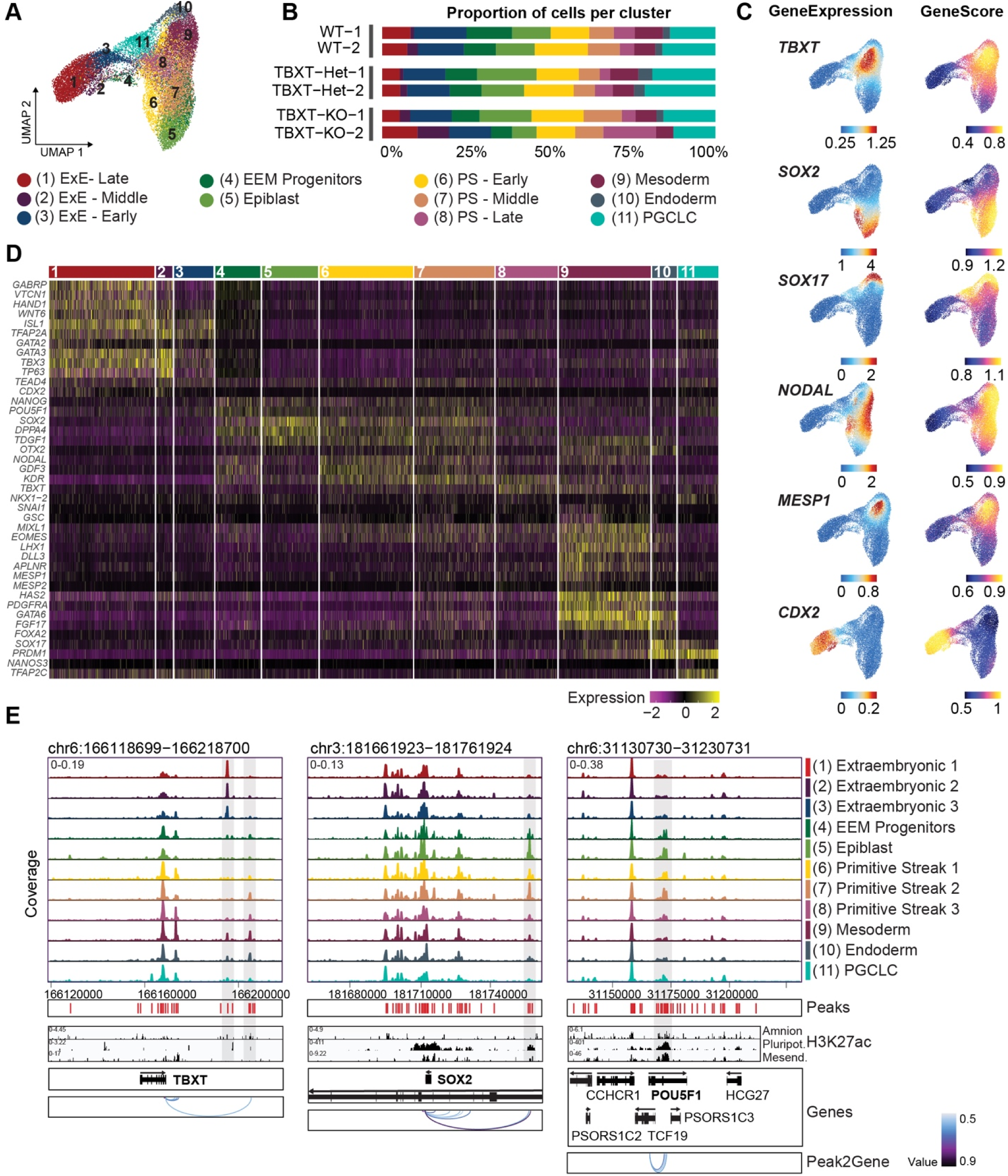
snRNA-seq reveals conserved lineage emergence in the absence of TBXT. (**A**) UMAP derived from snRNA-seq and snATAC-seq data depicting 11 clusters identified through Seurat analysis of snRNA-seq data after 48hr BMP exposure (nCells = 23,838, nReplicates = 2). (**B**) Bar Plot depicting the proportion of cells from each sample assigned to each cluster. (**C**) UMAP feature plots depicting GeneExpressionMatrix plots (derived from snRNA-seq) or GeneScoreMatrix plots for chromatin accessibility loci (derived from snATAC-seq) for key markers of gastrulation lineages. (**D**) Heatmap of snRNA-seq expression patterns for key markers of gastrulation lineages across clusters. (**E**) snATAC-seq accessibility tracks across clusters centered around TBXT, SOX2, and POU5F1 and aligned to detected peaks, H3K27ac tracks from published datasets, and Peak2Gene linkage predictions of regulatory connections between distal accessible regions (peaks) and nearby genes. Grey bars indicate peaks of interest.

To validate the conservation of this signaling network, we conducted IF for pSMAD1/5, WNT3a, NODAL, and the NODAL inhibitor LEFTY. The consistent localization of pSMAD1/5 along the colony periphery revealed that all genotypes maintain BMP4 signaling in this region (Fig. 1E, F). Additionally, NODAL was detected broadly and upregulated just interior toward the nascent mesoderm domain in gastruloids of all genotypes, and LEFTY was seen adjacent to this domain near the outer mesendoderm boundary. WNT3a protein expression was also maintained in the mesendoderm domain across all genotypes but was slightly decreased in this region of TBXT-KO gastruloids, suggesting the positive feedback loop between TBXT and canonical WNT signaling is disrupted in the TBXT-KO (Fig. 1E, F).

The uniformity in germ layer identity and distribution across genotypes suggests that cell fate specification is not dependent on TBXT dose during the early stages of human PS morphogenesis, and that cell fate specification and induction of morphogenetic movements are likely governed independently during *in vitro* gastrulation.

### Defining a genomic regulatory network underlying TBXT dosage

To gain a more comprehensive understanding of similarities to the malformations observed in *T/Bra* mutants observed *in vivo*, we sought to precisely define disparities in cell identity and the expression of migration regulators, including drivers of EMT, within gastruloids of each genotype. TBXT directly binds to the regulatory regions of key genes in mesoderm development and can influence chromatin accessibility (Faial et al., 2015; Koch et al., 2017), so we further hypothesized that TBXT expression level may influence chromatin accessibility at genes that are required for mesoderm maturation, and that altered accessibility may precede changes in protein level at the initial stages of mesoderm development (48-hour BMP4 exposure). To explore these possibilities, we conducted multiomic single nucleus RNA sequencing (snRNA-seq) and single nucleus Assay for Transposase-Accessible Chromatin (snATAC-seq) on gastruloids of each genotype after 48 hours of BMP4 treatment.

Our analysis of all 3 pooled genotypes yielded 11 clusters consisting of three extraembryonic cell populations (Clusters 1–3; “Extraembryonic-1–3”, extraembryonic progenitor cells (Cluster 4; “EEM Progenitors”), epiblast-like cells (Cluster 5; “Epiblast”), three primitive streak-like cell populations (Clusters 6–8; “PS-1–3“), nascent mesoderm-like cells (Cluster 9; “Mesoderm”), nascent endoderm-like cells (Cluster 10; “Endoderm”), and primordial germ cell-like cells (Cluster 11; “PGCLC”) (Figs. 2A-D, relationships between clusters at various resolutions illustrated in S3). In agreement with immunofluorescent data, we found there was not a significant difference in the proportion of cells from each genotype assigned to each cluster, supporting the notion that cell identity during early gastrulation was not significantly affected by the loss of TBXT.

Looking at these cluster identities in more depth, we observed that clusters 1–3 share an extraembryonic gene expression signature reflecting broad markers of both trophectoderm (TE) and amnion (*CDX2, GATA3, TFAP2A, HAND1, WNT6, GATA2*) (Fig. 2C-D). To distinguish these possibilities, we evaluated these clusters for key markers of trophectoderm, late-amnion, and early-amnion lineages. This analysis revealed a bias in all three clusters toward late-amnion (*GAPBRP, HEY1, HAND1, VTCN1, TPM1, IGFBP3, ANKS1A*) relative to embryonic clusters (Fig. S4), in agreement with published findings that primed hiPSCs are biased toward a late-amnion fate in the presence of BMP4 (Rostovskaya et al., 2022). This late-amnion gene expression signature was highest in cluster 1 and lowest in cluster 3, suggesting clusters 1–3 are likely distinguished by subtle variations in developmental timing with cluster 1 being a relatively more differentiated amnion and cluster 3 being nascent amnion. The companion snATAC-seq dataset revealed that clusters 1–3 have very similar chromatin accessibility and inferred peaks to both one another relative to all other clusters and to published H3K27AC accessibility data from primary amnion tissue (Bernstein et al., 2010), reinforcing their developmental similarity and amnion identity (Figs. 2E, S5D). These peaks are also enriched for GATA motifs, which are key regulators of extraembryonic cell fate (Fig. S5E). With these observations in mind, we designated “Extraembryonic-1” as “Extraembryonic-Late”, “Extraembryonic-2” as “Extraembryonic-Middle” and “Extraembryonic-3” as “Extraembryonic-Early”.

Following this developmental trajectory, cluster 4 shows a slight upregulation of extraembryonic markers (*GATA3, TFAP2A, HAND1, ISL1, TBX3*) relative to cluster 5, and cluster 5 exhibits canonical hallmarks of epiblast fate (*SOX2, POU5F1, NANOG, DPPA4*), including simultaneous motif enrichment of *SOX2*, *POU5F1*, and *NANOG* (Fig. 2C-D, S5E). Even with an apparent extraembryonic gene signature, cluster 4 maintains an epiblast-like gene signature overlapping with that of cluster 5. This gene expression pattern suggests that cluster 4 likely contains cells analogous to amnion progenitors leaving the PS, which we refer to as “Extraembryonic progenitors” (“EEM Progenitors”), while cluster 5 reflects epiblast-like cells (“Epiblast”).

The three PS clusters, clusters 6–8, share many elements of a PS gene signature, including *TBXT, MIXL1, and EOMES*, and are differentiated from one another by the progressive downregulation of epiblast markers such as *SOX2, TDGF1*, and *NODAL* (Fig. 2C-D). This pattern suggests that cluster 6 represents the least differentiated state between epiblast and mesoderm-like cells, followed by clusters 7 and then 8. Therefore, we designated “PS-1” (cluster 6) as “PS-Early”, “PS-2” (cluster 7) as “PS-Middle”, and “PS-3” (cluster 8) as “PS-Late.” Notably, clusters 4–8 share similar gene expression, chromatin accessibility, and peak distributions to one another, which reflects their recently shared epiblast-like origin and closely related identities (Fig. S5D). Analysis of differentially accessible regions (DARs) between clusters revealed a prospective SOX2 enhancer and the POU5F1 promoter that was uniquely accessible in the epiblast, extraembryonic progenitor, and early PS-like cell types (clusters 4–7) (Fig. 2E). These DARs correspond to H3K27ac ChIP-seq peaks in pluripotent HUES cells (Tsankov et al., 2015), reflective of these clusters’ pluripotency-related expression pattern and subsequent downregulation as differentiation progresses away from pluripotency.

Clusters 9 and 10, the nascent mesoderm and nascent endoderm-like clusters, respectively, share a mesendoderm-like gene expression signature (*GSC, MIXL1, EOMES, LHX1, HAS2, GATA6,* and *PDGFRA*). They are distinguished by the increased expression of *MESP1, MESP2, and APLNR* in mesoderm-like cells and *SOX17* and *PRDM1* in endoderm-like cells (Fig. 2C-D). In accordance, these two clusters are enriched for several motifs regulating mesendoderm specification, including GATA factors and FOXA2 (Fig. S5E). Chromatin accessibility near the *TBXT* promoter is increased in the PS and mesendoderm-like subclusters (clusters 6–10) and, interestingly, there appears to be a potential *TBXT* regulatory domain uniquely accessible in extraembryonic cell types (Fig. 2E). This peak is not linked to *TBXT* gene expression by Peak2Gene analysis, but it does correlate with the H3K27ac ChIP-seq profile of amnion tissue (Bernstein et al., 2010), suggesting that *TBXT* expression may be uniquely regulated in extraembryonic tissue relative to embryonic, a possibility that warrants further study. Finally, cluster 11 co-expresses *SOX17, PRDM1, NANOS3*, and *TFAP2C*, consistent with PGCLC identity (Fig. 2D).

### TBXT influences downstream gene expression in a dose-dependent manner

To understand why the loss of *T/Bra in vivo* most dramatically affects the morphogenesis of the mesoderm population, we assessed the nuanced differences that exist between the mesoderm populations of each genotype and how these may contribute to abnormal migration patterns observed later in development. We first compared differentially expressed (DE) genes between TBXT-Het vs. WT or TBXT-KO vs. WT within the mesoderm cluster (Fig. 3A-A’). This analysis identified 14 genes downregulated in the TBXT-Het compared with WT and 44 genes downregulated in the TBXT-KO compared with WT (Figs. 3B-D, S6A-B) (Adj. P < 0.05, log2FC < –0.25). Two general classes of DE genes emerged: genes that required a relatively binary threshold level of TBXT expression to fully activate or repress downstream expression and genes whose expression levels scaled with TBXT expression levels. Nine genes were statistically significantly downregulated in both the TBXT-Het and TBXT-KO compared to WT, putting them in the former category, including the non-canonical WNT pathway component *WNT5A* and BMP inhibitor *BMPER* (Fig. 3C-D, starred). In contrast, several genes including *WDPCP* and *TCF4* were significantly downregulated in the TBXT-KO compared to WT but had intermediate expression in the TBXT-Het (Fig. 3D, unstarred). GO Biological Pathway Enrichment for the 44 genes significantly downregulated in the TBXT-KO vs. WT, the majority of which also had reduced expression in the TBXT-Het, denoted “regulation of cell motility”, “locomotion”, and “PCP pathway involved in axis elongation,” reflecting the attenuated migratory phenotype and increased adhesiveness seen when TBXT expression is reduced *in vivo* (Wilson & Beddington, 1997)(Wilson et al., 1995) (Fig. 3E, Table S3). As anticipated, *in situ* hybridization confirmed decreased expression of the mesoderm marker *MESP1* and the WNT regulators *WNT5A* and *RSPO3* in TBXT-KO gastruloids relative to WT (Fig. 3F).

**Figure 3:**
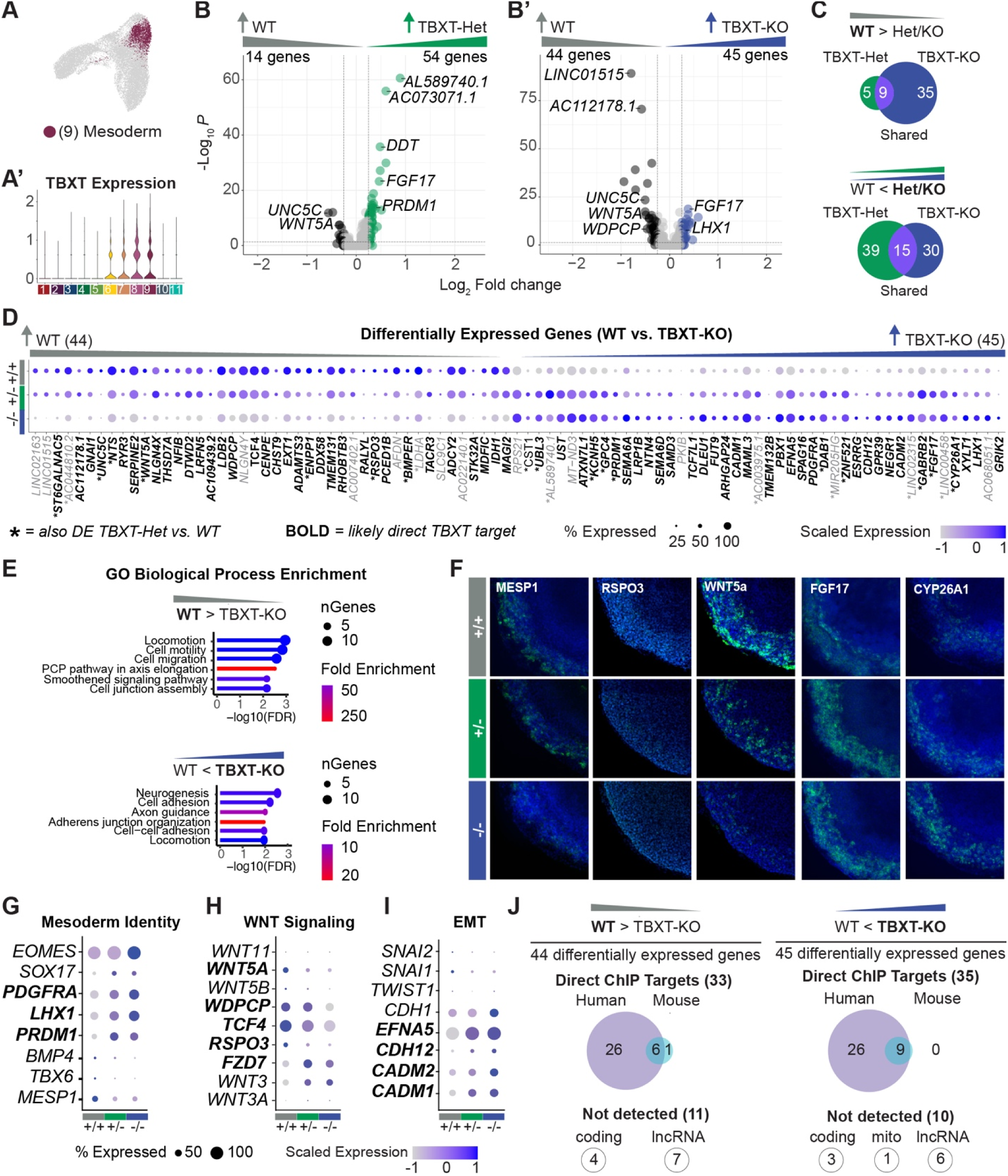
TBXT dose influences downstream gene expression in a dose-dependent manner. (**A**) UMAP highlighting the mesoderm cluster (Cluster “9”) and (**A’**) Violin plot showing the relative expression levels of TBXT in each cluster. (**B**) Volcano plot of differentially expressed genes between WT and TBXT-Het or (**B’**) WT and TBXT-KO after subsetting the mesoderm cluster (n = 3,212 cells, adj. p < 0.05, log2FC > 0.25) (**C**) Venn diagram of the number of DE genes identified in the TBXT-Het (green), TBXT-KO (Blue), or both TBXT-Het and TBXT-KO (purple) compared to WT. The top diagram includes genes downregulated in TBXT-Het and/or TBXT-KO compared to WT, while the bottom diagram includes genes upregulated in TBXT-Het and/or TBXT-KO compared to WT. (**D**) Dotplot of 89 DE genes between WT and TBXT-KO in order of log2FC. ‘*’ indicates genes that are also differentially expressed between WT to TBXT-Het, while the bolded black text indicates genes identified as likely direct targets of TBXT by comparison with ChIP-seq datasets. (**E**) Key results from ShinyGO GO Biological Process Analysis (FDR < 0.05, Pathway Size 2-2000 genes) (**F**) Multiplexed fluorescence *in situ* hybridization for *MESP1*, *RSPO3, WNT5A, FGF17*, and *CYP26A1* in gastruloids at 48hr. *In situ* hybridizations were repeated for >=3 gastruloids of each genotype. (**G-I**) Dotplots showing gene expression trends for key genes related to mesoderm identity (**G**), WNT signaling (**H**), and EMT (**I**). (**J**) Venn diagrams summarizing DE genes detected in corresponding ChIP-seq datasets.

On the other hand, we identified 54 genes upregulated in the TBXT-Het compared to WT and 45 genes upregulated in the TBXT-KO compared to WT (Fig. 3B-D, S6A) (Adj. P < 0.05, log2FC > 0.25). Of these, 15 genes overlapped, including the posterior morphogens *CYP26A1* and *FGF17* and the endoderm marker *PRDM1* (Fig. 3C-D, starred). Genes that demonstrated expression patterns that scaled with TBXT dose included migration and adhesion ligand *EFNA5*, cell adhesion molecules *CADM1*, *CADM2,* and *CDH12*, and endoderm marker *LHX1* (Fig. 3D). GO enrichment of the 45 genes significantly upregulated in the TBXT-KO denoted “cell-cell adhesion” and “adherens junction organization,” reflective of a persistent epithelial character and impaired migration in the absence or reduction of TBXT (Fig. 3E, Table S3). In addition, several terms reflective of neurogenesis emerged. These terms are largely driven by genes involved in axon guidance or migration such as *NTN4*, *SEMA6A*, and *EFNA5*, so this may reflect a neural bias or a misregulation of genes involved in the migratory mesenchymal phenotype. We conducted *in situ* hybridization for *FGF17* and *CYP26A1* to validate these results and observed increased expression of both transcripts in TBXT-KO gastruloids relative to the WT (Fig. 3F).

Overall, the level of expression of differentially expressed genes in the TBXT-Het was always similar to the TBXT-KO, similar to the WT, or an intermediate between the two. This pattern suggests that while some genes respond to TBXT dose in a binary manner where WT expression levels are needed for downstream expression, more often the expression level of TBXT is closely linked to the expression level of its downstream targets.

### TBXT dose influences the expression profile within the mesodermal population

While the presence and spatial distribution of broad mesoderm markers were not affected by varying TBXT expression, closer examination revealed that the gene regulatory network controlling early mesendoderm identity itself is subtly influenced by TBXT dose. The canonical endoderm and PGCLC marker, *PRDM1*, was significantly upregulated in both the TBXT-Het and the TBXT-KO relative to WT, suggesting its expression is sensitive to TBXT dose (Fig. 3D, G, S6A, Table S4). *LHX1* and *PDGFRA* had a similar expression pattern as *PRDM1*, with intermediate expression levels in the TBXT-Het and significant upregulation in the TBXT-KO compared to WT. *SOX17* and *EOMES* also followed a similar pattern, with slightly elevated expression in the TBXT-KO relative to the WT. In contrast, mesoderm markers *MESP1*, *TBX6,* and *BMP4* were slightly higher in the WT compared to the KO (Fig. 3G, S6C). Canonical WNT pathway components *RSPO3*, *WDPCP*, and *TCF4* also displayed significantly graded expression, with the highest expression in WT and the lowest in the TBXT-KO (Fig. 3D, H, S6D). *WNT3A* itself was very sparsely detected in the nascent mesoderm population via snRNA-seq, however, the significant upregulation of several canonical WNT pathway components in WT agrees with the WNT3A expression pattern observed via IF (Fig. 1D-F, 3H).

Notably, several direct targets of TBXT that are implicated in osteogenesis were significantly upregulated in WT, including *BMPER*, *GNAI2, ENPP1, THSD7A,* and *ADAMTS3* (Fig. S6C). This is relevant because *Tbxt*^+/-^ mutant mice frequently display skeletal malformations later in development including rib fusions, osteochondrodysplasia, and brachydactyly (Grüneberg, 1958; Herrmann et al., 1990; Wilson et al., 1993). In addition, *MAML3*, which amplifies transcription of *HES1* to drive oscillations in somitogenesis, was increased in the TBXT-KO gastruloids. This expression change is potentially related to somitic fusions and subsequent rib fusions seen in *Tbxt*^+/-^ and *Tbxt*^-/-^ animal models later in development (William et al., 2007; Wu et al., 2002). Finally, the BMP inhibitor *BMPER* was significantly downregulated in both TBXT-Het and TBXT-KO compared to WT. Because BMP4 signaling activates TBXT expression and TBXT directly activates BMPER expression, decreased *BMPER* expression in the TBXT-KO could reflect a compensatory pathway to rescue TBXT expression levels in TBXT-Het or TBXT-KO.

These gene expression trends suggest that while reduced TBXT expression does not preclude cells from forming mesoderm, it does result in an endoderm-biased gene expression profile in this nascent mesoderm and influence the gene regulatory networks related to downstream axial elongation and patterning.

### TBXT dose influences the expression of genes that modulate EMT

In addition to genes related to endoderm identity, we observed several gene expression patterns reflective of impaired EMT in our mutant gastruloids. For example, *CDH1* was upregulated in the TBXT-KO mesoderm, intermediate in the TBXT-Het mesoderm, and downregulated in the WT mesoderm (Fig. 3I). Cell adhesion molecules *CADM1, CADM2*, *CDH12*, and *EFNA5* shared this expression pattern, suggesting that the mutant cell lines retain an epithelial character, unlike their WT counterparts. Indeed, *SNAI1* and *SNAI2*, canonical regulators of EMT, both had the highest expression in the WT mesoderm, although they were detected in very few cells. The upregulation of adhesion molecules and downregulation of SNAI family proteins observed in TBXT-Het and TBXT-KO suggests that while the WT population is actively undergoing EMT, this process is impaired in the mutant gastruloids.

### The majority of differentially expressed genes are direct targets of TBXT

To better isolate the key components of the gene regulatory network underlying TBXT dose-responsive gene expression, we sought to identify which differentially expressed genes are also likely direct targets of TBXT based on promoter binding proximity. To do this, we leveraged four existing TBXT chromatin immunoprecipitation with sequencing (ChIP-seq) datasets from human and mouse embryonic stem cells (hESC/mESCs) grown *in vitro*. The first two datasets were derived from hESCs differentiated into monolayer TBXT+ cell populations using either activin (endoderm-biased) or BMP4 (mesoderm-biased) protocols (Faial et al., 2015). The third dataset utilized a mesendoderm-biased hESC population driven by TGF-β and WNT signaling (Tsankov et al., 2015). The final study used an Activin-A mediated protocol to drive mESC embryoid bodies to a Tbxt+ PS fate (Lolas et al., 2014).

This comparison revealed that the vast majority of the genes that are differentially expressed between the different genotypes within the mesoderm cluster have TBXT binding sites adjacent to their promoters and are therefore likely direct targets of TBXT. Specifically, 33 of the 44 differentially expressed genes downregulated in TBXT-KO compared to WT had proximal TBXT binding sites in at least one ChIP-seq dataset (Fig. 3D (bold), 3J, S7). Six of these 33 genes were detected in both human and mouse ChIP-seq datasets (*BMPER, DTWD2, EXT1, MDFIC, ENPP1,* and *RSPO3*), suggesting they play an evolutionarily conserved function in early development. The remaining 26 genes were specifically identified in the human ChIP-seq datasets. One gene, *PCED1B*, was detected as a potential direct target of TBXT in the mouse datasets but not in any of the human datasets. Seven of the 33 differentially expressed genes identified in the ChIP-seq datasets were significantly downregulated in both the TBXT-Het and TBXT-KO colonies compared to WT, including *BMPER*, *WNT5A*, and *UNC5C,* suggesting these genes are particularly sensitive to reduced TBXT expression. The 11 differentially expressed genes that were not identified as likely direct targets of TBXT included 4 protein-coding genes (*LDHA, AFDN, NLGN4Y,* and *SLC9C1*) and 7 long non-coding RNAs.

Thirty-five out of the 45 genes upregulated in the TBXT-KO compared to WT were found to have proximal TBXT binding sites in at least one ChIP-seq dataset (log2FC > 0.25, Adj. P < 0.05) (Fig. 3D (bold), 3J, S7). Of these, 9 were identified as potential direct targets of TBXT in both human and mouse datasets (*FGF17, LHX1, SAMD3, UBL3, CADM1, EFNA5, MAML3, SEMA6A, and TCF7L1).* The remaining 26 genes were identified specifically in the human ChIP-seq datasets. Nine of the 35 potential direct targets, including *FGF17, CYP26A1,* and *PRDM1,* were significantly upregulated in both TBXT-Het and TBXT-KO colonies compared to WT, reflecting a more pronounced dosage sensitivity (Fig. S7 (bold)). Ten differentially expressed genes were not identified in any of the ChIP-seq datasets, of which 3 were detected in coding regions (*CST1*, *PKIB*, and *RPS21*), 1 was mitochondrial, and the remaining 6 were long non-coding RNAs.

Taken together, these results indicate that the majority of genes that display TBXT dosage sensitivity have proximal binding sites for TBXT, and therefore their transcription is likely directly modulated by TBXT.

### TBXT does not significantly impact chromatin accessibility in nascent mesoderm

It has previously been shown that TBXT plays a role in the deposition of H3K27ac at target genes during hematopoietic and endothelial development to alter transcription (Beisaw et al., 2018; Chen et al., 2023) and that TBXT is essential for remodeling chromatin during NMP development (Koch et al., 2017). We were interested in determining whether TBXT similarly modulates chromatin accessibility during early PS morphogenesis. However, in our gastruloid mesoderm population, very few genomic regions were differentially accessible between either the TBXT-Het and WT or TBXT-KO and WT conditions (26 and 6 genes, respectively; FDR < 0.1, log2FC > 0.5) (Fig. S8A-C, Table S4). While it is true that TBXT-Het does reflect a higher number of differentially accessible regions (DARs) than TBXT-KO when compared to WT, the log2FC values of these DARs are very close to the significance cutoff and several are microRNAs. The 6 DARs identified in the TBXT-KO included adhesion protein *MDGA2*, vitamin D metabolizing enzyme *CYP2R1*, pluripotency-related gene *EYS*, histone components *HIST1H4L*, *HIST1H1B*, and *HIST1H3I*, and several long non-coding RNAs. Likewise, very few peaks were differentially accessible between the two conditions (18 peaks between WT and TBXT-Het and 0 peaks between WT and TBXT-KO, respectively; FDR < 0.1, log2FC > 0.5), and no motifs were significantly enriched within these peaks, as was true for all clusters (Table S4). We observed slight variation in peak height at genes demonstrating differential expression such as *WNT5A, RSPO3*, and *LHX1*, although these variations did not reach statistical significance (Fig. S8D). These results suggest that at this early stage of mesoderm specification in gastruloids, TBXT expression does not contribute to mesodermal patterning by influencing chromatin accessibility.

### CellChat reveals TBXT-driven regulation of cell-cell adhesions and ncWNT signaling across and within clusters

Morphogenesis frequently involves paracrine and juxtracrine signaling between different cell populations. For example, mesodermal cells in contact with the epiblast and visceral endoderm have distinct protrusions whereas mesoderm cells in contact with other mesoderm cells appear smoother, reflective of distinct cell responses to specific migratory guidance cues (Saykali et al., 2019). Therefore, we questioned whether TBXT-dose-dependent changes in cell behavior are restricted to the mesoderm population or if they may be influenced by adjacent cell types. To begin to address this question, we turned to the software CellChat, which analyzes the expression of ligand-receptor pairs within and across clusters to predict patterns in cell-cell communication.

First, we investigated how TBXT expression influences broad patterns in pathway activation by assessing which pathways have the largest changes in signals between or within different cell types, termed ‘information flow,’ when comparing WT, TBXT-Het, and TBXT-KO snRNA-seq data across all 11 clusters. This analysis revealed several pathways that have varying levels of information flow, and we focused on the three in which the WT and TBXT-KO have distinctly different patterns from one another (Fig. 4A). The first two pathways, cell adhesions (CADM) and junctional adhesions (JAM), were predicted to have higher information flow in the TBXT-Het and TBXT-KO gastruloids relative to WT, while the third pathway, non-canonical WNT (ncWNT) signaling, was predicted to have higher information flow in WT and TBXT-Het gastruloids relative to TBXT-KO.

**Figure 4:**
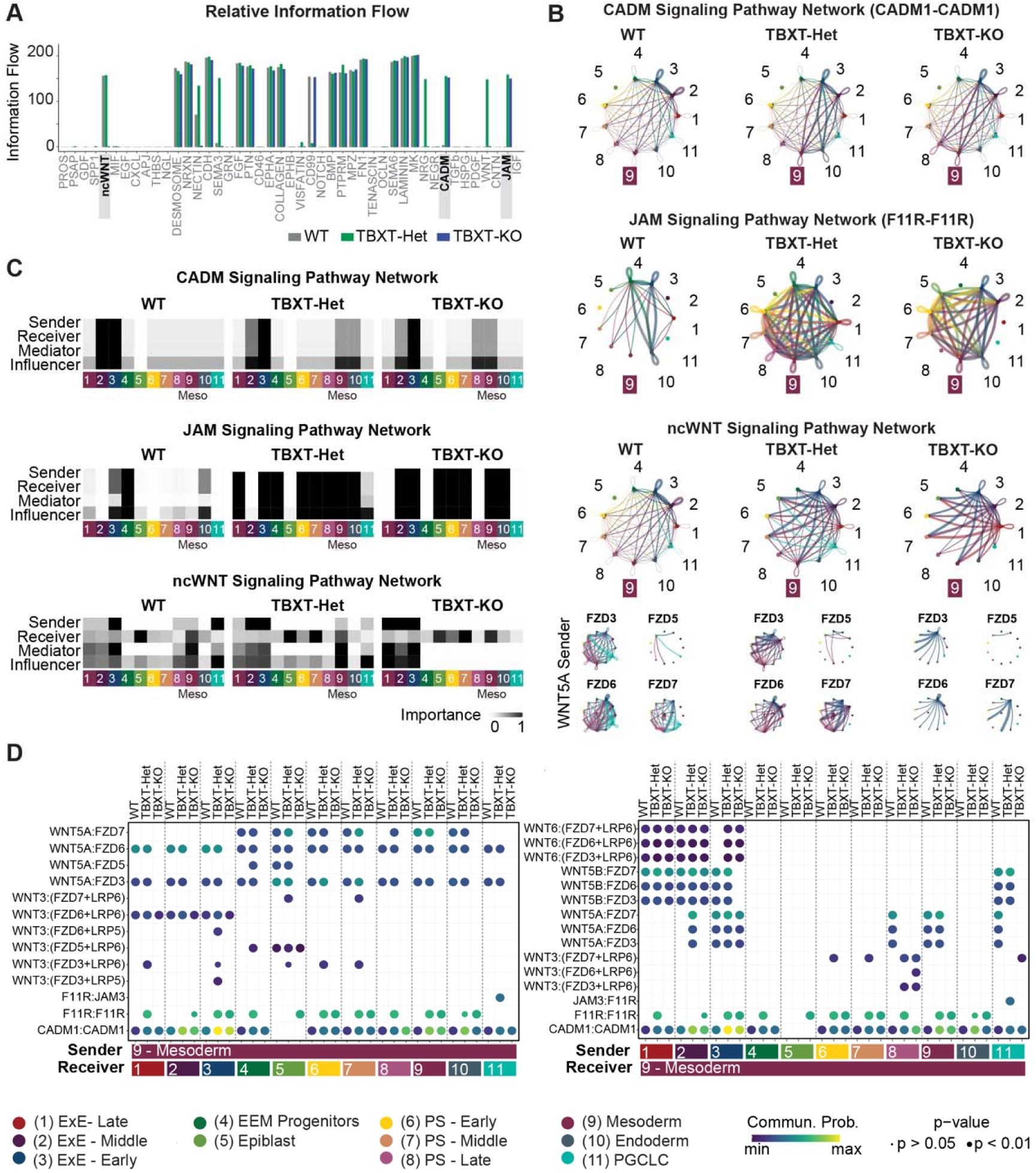
CellChat reveals TBXT-driven regulation of cell-cell adhesions and ncWNT signaling across and within clusters. (**A**) Bar plot comparing information flow (A. U.) of key pathways between WT, TBXT-Het, and TBXT-KO. (**B**) Circle plots visualizing communication between or within clusters for the CADM, JAM, and ncWNT signaling pathways. Numbers correlate with cluster identity, while line thickness corresponds to the strength of predicted communication. “9” indicates the mesoderm cluster. Small circle plots under the “ncWNT Signaling Pathway” header depict interactions between WNT5A and designated FZD receptors. (**C**) Heatmap showing the communication dynamics for each genotype for the CADM, JAM, and ncWNT signaling pathways. (**D**) Bubble plot comparing predicted ligand-receptor interactions for CADM, JAM, WNT, and ncWNT pathways. Only ligand-receptor interactions with variability in communication probability are shown.

To understand how this expression pattern affects interactions across clusters, we looked at predicted ligand-receptor interactions of each of these three pathways within and between each cluster for all three genotypes. Analysis of CADM pathway and JAM pathway communication across and with clusters revealed these hits were primarily driven by varying expression levels of *CADM1* and *F11R*, respectively, although the JAM pathway was also influenced by *JAM3* expression (Fig. S9A). We then visualized ligand-receptor interactions between and within clusters using circle plots, where the presence and thickness of a line correlate with the degree of predicted communication within or between clusters. An overall increase in CADM signaling was evident in TBXT-Het and TBXT-KO gastruloids relative to WT, and this seemed to be largely due to increased communication within the PS-Late and mesoderm clusters (Fig. 4B). Similarly, in TBXT-Het and TBXT-KO gastruloids, JAM signaling appeared exaggerated in PS and mesoderm clusters whereas these signaling dynamics were largely lost in WT. These trends suggest that unlike the WT, the TBXT-Het and TBXT-KO colonies are maintaining their cell-cell adhesions and junctional adhesions, and these differences are uniquely apparent in cell types expected to undergo EMT such as PS-Late and Mesoderm.

Within the ncWNT pathway, the TBXT-Het and TBXT-KO gastruloids had reduced ligand-receptor interactions compared to WT both between clusters but also within clusters, with the degree of self-regulation changing most notably within the three PS clusters and the mesoderm cluster (Fig. 4B). This variation in ncWNT signaling across genotypes was driven by the expression patterns of *WNT5A*, *WNT5B*, and several *FZD* receptors (Fig. 4B, S9A-C). WNT5A is a direct target of TBXT and a crucial component of both the non-canonical WNT/Planar Cell Polarity (PCP) pathway, which controls many aspects of directed cell migration and convergent extension, and the canonical WNT pathway, which is critical for sustained mesoderm development (Dunty et al., 2008; Kikuchi et al., 2012; Yamaguchi et al., 1999).

To better understand how CADM, JAM, and ncWNT pathways are influenced by TBXT dose, we next explored the extent to which different clusters operate as senders, receivers, mediators, and influencers of ligands and receptors in these pathways. Mediators serve as gatekeepers to control cell communication between any two groups, while influencers are predicted to control information flow more generally (Jin et al., 2021). For both the CADM and JAM pathways, the TBXT-Het and TBXT-KO PS-late and mesoderm clusters increased their ability to serve as senders, receivers, mediators, or influencers relative to WT (Fig. 4C). These trends likely reflected genotype-dependent patterns in *CADM1* and *F11R* expression within these specific clusters. In contrast, the PS-late, mesoderm, and PGCLC clusters largely lost their ability to operate as senders, mediators, and influencers of the ncWNT pathway, but they maintained their ability to serve as receivers (Fig. 4C). These patterns suggest that variability in ncWNT pathway information flow is likely more highly dependent on varying levels of *WNT5A* or *WNT5B* expression (senders) rather than *FZD* expression (receivers).

To clarify how specific ligands or receivers modulated predicted information flow in the ncWNT pathway, we looked at how these gastruloid communication patterns were affected by the expression of specific ligand-receptor pairs sent or received by the mesoderm cluster (Fig. 4D). We found that WNT5A signals sent from the mesoderm to FZD receptors across all other clusters, including the mesoderm itself, were largely lost in the TBXT-KO but maintained in WT. WNT5B, however, was not detected as a major ligand sent from the mesoderm. Reciprocally, ncWNT signals such as WNT5A and WNT5B sent from extraembryonic-early, PS-late, and PGCLC clusters to the mesoderm cluster showed increased signal in the WT, likely reflecting an increase in *WNT5A* expression in all TBXT-expressing clusters. This analysis also revealed that WNT5B is most impactful in extraembryonic and PGCLC clusters, but relatively negligible in PS and mesendodermal clusters.

Taken together, our analyses directly reflect the persistence of cell adhesions and junctional adhesions in the TBXT-Het and TBXT-KO PS-late and mesoderm clusters, suggesting that these cells do not readily acquire a mesenchymal identity, a key requirement for motility and subsequent PS morphogenesis. The results additionally reflect the downregulation of *WNT5A* in the TBXT-Het and TBXT-KO compared to WT and identify the non-canonical WNT signaling pathway as a key TBXT-dependent regulator of nascent mesoderm development.

### TBXT dose influences the persistence of cell-cell adhesions

Gastruloid colonies develop a very dense mesoderm layer that prohibits clear visualization of cell morphology and junctions. Therefore, to understand how cell-cell adhesions and EMT are affected by TBXT dose, we turned to an alternate protocol that induces early mesoderm in a monolayer using StemCell Technology’s Mesoderm Induction Media (MIM). In WT cells, this media induces TBXT expression in >90% of cells within 48 hours, thereby recapitulating the initial stages of nascent mesoderm specification (Fig. 5A-B, S10A). This early mesoderm population maintains consistent EOMES expression regardless of TBXT genotype with expression peaking at 48hrs, reinforcing the shared nascent mesoderm identity across genotypes (Fig. S10B). At 48 hours we observe a TBX6+ mesoderm population in WT and TBXT-Het and a reduced but present TBX6+ population in TBXT-KO. This is in agreement with our snRNA-seq data and the established role of TBX6 as a direct target of TBXT in the PS and later in the paraxial mesoderm. (Fig. 3G, S10C). We differentiated cells of each genotype using MIM for either 0hr, 24hr, 48hr, or 72hr, and assessed cell morphology and protein expression via IF.

**Figure 5:**
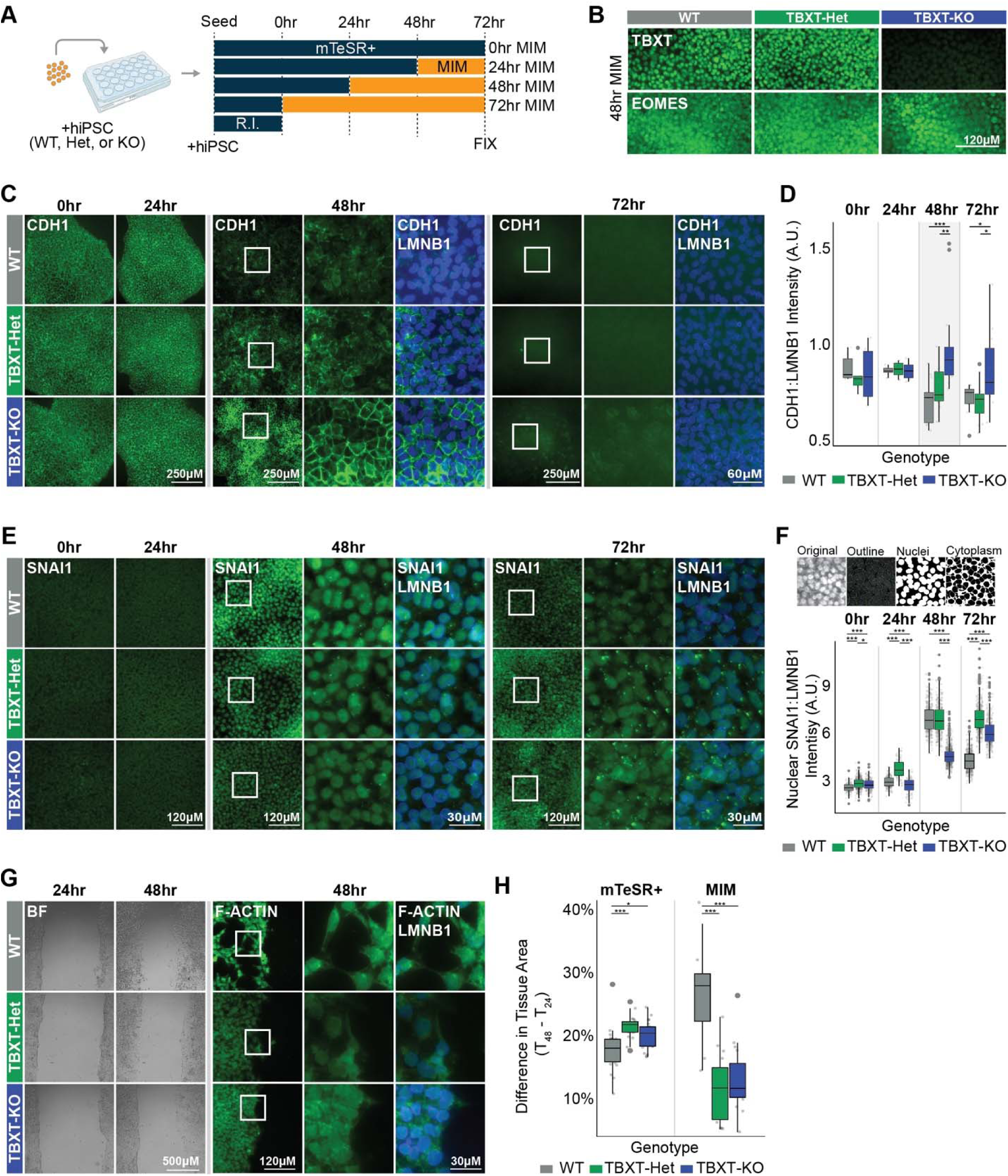
TBXT dose directly impacts the timing of EMT to permit migration. (**A**) Schematic of the experimental setup. MIM = Mesoderm Induction Media. “R.I.” indicates ROCK inhibitor. (**B**) Immunofluorescence for TBXT or EOMES expression in each genotype after 48hr MIM exposure. (**C**) Immunofluorescence for CDH1 or (**E**) SNAI at 0hr, 24hr, 48, or 72hr MIM exposure. (**D**) Quantification of the ratio between CDH1 and LMNB1 intensity. n >= 6 wells/genotype (**F**) Schematic showing segmentation of nuclei and cytoplasm and the quantification of the ratio between SNAI and LMNB1 specifically within cell nuclei. WT = 1,228 cells; TBXT-Het = 1,423 cells TBXT-KO = 1,373 cells. n = 3 images/genotype/timepoint. (**G**) Brightfield images of the scratch assay at 24hrs or 48hrs MIM exposure (0hr or 24hr relative to scratch). Immunofluorescence for F-ACTIN at the edge of a scratch wound at 48hrs (24hr relative to scratch). (**H**) Quantification of the difference in tissue area between 48hr and 24hr MIM exposure (n = mTeSR+: WT= 23, TBXT-Het = 19, TBXT-KO = 21. MIM: WT = 8, TBXT-Het = 15, TBXT-KO = 16).

Varying the timing of MIM exposure revealed a robust and stepwise delay in the timing of the downregulation of CDH1, a canonical marker of the epithelial state, across genotypes (Fig. 5C-D). At 24 hours of MIM treatment, all genotypes displayed robust junctional CDH1 expression. However, by 48 hours, CDH1 was largely eliminated in WT cells, TBXT-Het cells had variable expression, and TBXT-KO cells maintained robust junctional expression, in agreement with our snRNA-seq data. By 72 hours, TBXT-Het cells had lost CDH1 expression to mirror WT cells, whereas CDH1 was reduced but still detectable in TBXT-KO cells. This expression pattern demonstrates that TBXT dose is directly correlated to the temporal downregulation of CDH1 in the nascent mesoderm, which in turn correlates with stunted EMT progression.

To better understand how TBXT influences EMT progression more broadly, we looked at the distribution of SNAI1, an established marker of EMT and inhibitor of CDH1 (Muqbil et al., 2014; Xu et al., 2019), across the same 72hr timespan. As previously indicated, we detected higher *SNAI1* RNA expression in WT compared to TBXT-Het and TBXT-KO via snRNA-seq (Fig. 3I). IF revealed that WT and TBXT-Het had a significantly higher level of nuclear SNAI1 relative to TBXT-KO at the 48hr timepoint (Fig. 5E-F). At 72hr this expression was beginning to drop off in the WT but was maintained in the TBXT-Het and TBXT-KO, revealing an inversed correlation to CDH1 expression. Furthermore, we also observed increased protein expression of other epithelial markers including junctional β-catenin and ZO1 in the TBXT-KO cells relative to the WT after 48hr MIM, in accordance with TBXT dose-dependently augmenting the progression of EMT (Fig. S11A).

EMT progression and mesenchymal cell motility are linked to alterations in extracellular matrix composition. In particular, the deposition of the basement membrane protein and SNAI1-target fibronectin (FN1) reflects the acquisition of a mesenchymal phenotype. We observed decreased deposition of basement membrane component FN1 in our TBXT-KO relative to TBXT-Het and WT, reflecting maintained cell adhesions and decreased motility in the absence of TBXT (Fig. S11A).

Next, we utilized the scratch wound assay to visualize the migration patterns of cells of each genotype and quantify how TBXT-dose-dependent changes in EMT-related protein expression correlate with cell migration kinetics. Comparing the difference in the percent of occupied tissue area between the starting time point (24hr MIM) and ending time point (48hr MIM) across genotypes revealed that the WT population migrated significantly farther into the wound space than the TBXT-Het or TBXT-KO populations (Fig. 5G-H). WT cells were noticeably larger and much more motile both as a group and independently. Staining with F-Actin revealed long protrusions in WT cells, reflective of their motile mesenchymal character. TBXT-KO cells, in contrast, remained tightly packed and epithelial, and cell movement appeared to be driven by overconfluence more so than directed cell movement. A small fraction of TBXT-Het cells acquired a similar mesenchymal phenotype to WT, but the vast majority appeared epithelial and non-migratory. Therefore, increased SNAI1 and decreased CDH1 expression seen in WT cells at the 48-hour MIM timepoint reflect the acquisition of a TBXT-dependent migratory mesenchymal phenotype.

The expression patterns of CDH1, β –catenin, ZO1, SNAI1, and FN1 in addition to the distinct migratory dynamics seen across genotypes indicate that TBXT plays a crucial role in regulating the temporal component of EMT, where TBXT dose-dependently promotes the reduction of CDH1, nuclear localization of SNAI, and subsequent acquisition of a mesenchymal phenotype.

## DISCUSSION

Our study demonstrates that the acquisition of mesodermal identity can be decoupled from the acquisition of a mesenchymal character, and TBXT is required for the latter in a dose-dependent manner. After 48 hours of BMP4 exposure, the mesoderm cluster of our gastruloids exhibit TBXT-dose dependent changes in gene expression related to mesendoderm identity, both non-canonical and canonical WNT signaling, and EMT, and the majority of these differentially expressed genes are likely directly regulated by TBXT as evidenced by comparisons to existing ChIP-seq datasets. CellChat analysis highlighted the JAM, CADM, and ncWNT pathways as uniquely regulated between WT and TBXT-KO colonies and showed that the mesoderm and PS-late clusters play key roles in the regulation of these pathways. We then utilized a 2D monolayer culture system to demonstrate that TBXT dose directly correlates with the temporal downregulation of genes related to cell adhesions, the upregulation of drivers of EMT, and the subsequent acquisition of a mesenchymal phenotype.

These findings have interesting implications for our understanding of how TBXT influences embryonic patterning at the earliest stages of gastrulation, even before the commencement of posterior trunk development. As opposed to NMP populations which require TBXT for mesoderm differentiation (Gouti et al., 2017; Koch et al., 2017), we demonstrated that early PS populations do not require TBXT for the initial establishment of a relatively anterior mesoderm identity. This result is in agreement with studies showing that another T-box factor, EOMES, primarily drives mesoderm differentiation at this stage of development (Schüle et al., 2023). However, we also discovered that TBXT is not dispensable at this early stage, as its expression directly influences the timing of EMT and therefore the ability of the nascent mesoderm cells to properly acquire a migratory mesenchymal character. We believe that this impaired EMT observed *in vitro* is reflected in the “pile-up” phenotype observed *in vivo,* as cells accumulate in the PS when *T/Bra* expression is lost.

Somewhat surprisingly, we did not observe large-scale changes in chromatin accessibility across TBXT genotypes, despite detecting variation in gene expression. While this may be specific to our *in vitro* model system, it likely better reflects the relatively early time point modeled by the gastruloids. At 48 hours mesoderm is just beginning to emerge, and it is possible that with continued differentiation over an additional 24-48 hours, the mesoderm identity would advance, and chromatin remodeling would be more readily apparent. It is also possible that TBXT does not directly remodel chromatin during early gastrulation and might instead be interacting with histone-modifying complexes.

Throughout this study we focused on cell populations analogous to those leaving the PS, however, extraembryonic tissues were also affected by the loss of TBXT. Like in the PS and mesoderm, CellChat specifically identified alterations in signaling within the JAM, CADM, and ncWNT pathways in extraembryonic populations. TBXT-KO embryos die from asphyxiation due to impaired allantois development, and so it will be interesting to explore the extent to which this is driven by analogous defects in EMT that limit extraembryonic cell migration.

Overall, this study clarifies the role of TBXT during early PS development and sheds light on its ability to promote the temporal progression of EMT in a dose-dependent manner. While TBXT-KO cells do ultimately downregulate CDH1 and acquire a motile phenotype, this transition occurs considerably later than in WT cells, demonstrating that the correct timing of EMT is critical for proper morphogenesis. In addition, this study decouples EMT progression from initial mesoderm specification in the PS and complements *in vivo* studies to both improve our core understanding of vertebrate mesoderm development and identify nuances of human development. This understanding will help us to effectively design directed differentiation strategies that incorporate both EMT and cell fate acquisition, in addition to improving our foundational understanding of human embryogenesis.

## METHODS

### Cell Lines

All work with human induced pluripotent stem cells (hiPSCs) was approved by the University of California, San Francisco Human Gamete, Embryo, and Stem Cell Research (GESCR) Committee. Human iPS cells harboring genome-edited indel mutations for *TBXT* (TBXT-Het, TBXT-KO) were generated for this study and derived from the Allen Institute WTC11-LaminB parental cell line (AICS-0013 cl.210). The WT line was derived from a WTC11-LaminB subclone that was exposed to the TBXT sgRNA but remained unedited. All cell lines were karyotyped by Cell Line Genetics and reported to be karyotypically normal. Additionally, all cell lines tested negative for mycoplasma using a MycoAlert Mycoplasma Detection Kit (Lonza).

### Maintenance of iPS Cells

Human iPSCs were cultured on growth factor-reduced (GFR) Matrigel (Corning Life Sciences) and fed at minimum every other day with mTeSR-Plus medium (STEMCELL Technologies) (Ludwig et al., 2006). Cells were passaged by dissociation with Accutase (STEMCELL Technologies) and re-seeded in mTeSR-Plus medium supplemented with the small molecule Rho-associated coiled-coil kinase (ROCK) inhibitor Y276932 (10 μM; Selleckchem) (Park et al., 2015) at a seeding density of 12,000 cells per cm^2^. After 24 hours, cells were maintained in mTeSR-Plus media until 80% confluent.

### TBXT Allelic Series Generation

To generate the TBXT allelic series we first lipofected WTC11-LaminB cells (AICS-0013 cl.210) with 125ng sgRNA and 500ng Cas9 protein according to the Lipofectamine Stem Transfection Reagent Protocol (Invitrogen). The human TBXT sgRNA (CAGAGCGCGAACTGCGCGTG) was a gift from Jacob Hanna (Addgene plasmid #59726; http://n2t.net/addgene:59726; RRID: Addgene_59726) and targeted the first exon of the TBXT gene. After recovery for 48 hours in mTeSR-Plus supplemented with ROCK inhibitor, lipofected cells were dissociated using Accutase and passaged into a GFR-Matrigel coated 10cm dish, where they were expanded for 24hr in mTeSR-Plus with ROCK inhibitor. Media was replaced with mTeSR-Plus without ROCK inhibitor and cells continued to grow another 2-4 days before the manual selection of 20-30 single colonies into individual wells of a 96-well plate. After the expansion of the clonal populations for 5-10 days, cells were passaged into a new 96-well plate at a 1:5 dilution ratio in mTeSR-Plus supplemented with ROCK inhibitor, and the remaining cells were used for genotyping. For screening of *TBXT* exon 1 non-homologous end-joining (NHEJ) mutations, DNA was isolated using QuickExtract DNA lysis solution (Epicentre #QE0905T), and genomic DNA flanking the targeted sequence was amplified by PCR (For1: gaaggtggatctcaggtagcgagtctgg and Rev1: cagcaggaaggagtacatggcgttgg) and sequenced using Rev1. Synthego ICE analysis was employed to quantify editing efficiency and identify clones with heterozygous (45-55% KO score) or homozygous null (>90% KO score) mutations (Table S1). To eliminate the possibility of a heterozygous line being a mixed population of wildtype and homozygous null alleles, 8-12 subclones of the prospective heterozygous clones were isolated and Sanger sequenced as before, and all subclones were confirmed to contain identical genotypes (Table S1). After sequencing confirmation of respective genotypes, karyotypically normal cells from each hiPSC line were expanded for subsequent studies.

### PDMS stamp fabrication

Standard photolithography methods were used to fabricate a master template, which was provided as a gift from PengFei Zhang and the Abate lab at the University of California, San Francisco (Alom Ruiz & Chen, 2007; Minn et al., 2020; Théry & Piel, 2009). The photoresist master was then coated with a layer of chlorotrimethylsilane in the vacuum for 30 min. Polydimethylsiloxane (PDMS) and its curing agent, Sylgard 184, (Dow Corning, Midland, MI) were mixed in a 10:1 ratio, degassed, poured over the top of the master, and cured at 60°C overnight, after which the PDMS layer was peeled off to be used as a stamp in micro-contact printing.

### Microcontact Printing

PDMS stamps (each containing 12 x 1000uM circles) were sterilized by washing in a 70% ethanol solution and dried in a laminar flow hood. Growth factor reduced Matrigel (Corning) was diluted in DMEM/F-12 (Gibco) at 1:100 dilution and incubated on the stamps to cover the entire surface of the feature side at 37°C for 1 hour. The Matrigel solution was then aspirated off the stamps, which were air-dried. Using tweezers, the Matrigel-coated surface of stamps was brought in contact with glass or plastic substrate, usually a glass 24-well plate or removable 3-chamber slide (Ibidi), and incubated on the substrate for 1hr at 37°C. The stamps were then removed and rinsed in ethanol for future use. Matrigel-printed substrates were incubated with 1% Bovine Serum Albumin (Sigma Aldrich) in DPBS-/-at room temperature for 1 hour before being stored in DPBS-/-solution at 4°C for up to 2 weeks.

### Confined 2D Gastruloid Differentiation

hiPSCs were dissociated with Accutase and resuspended in mTeSR-Plus supplemented with ROCK inhibitor, as previously described. Cells were then seeded onto a stamped well at a concentration of approximately 750 cells/mm^2^. Cells were incubated at 37°C for 3 hours before the well was rinsed 1x with DPBS and given fresh mTeSR-Plus supplemented with ROCK inhibitor. Approximately 24 hours post-seeding, media was exchanged for mTeSR-Plus. After another 24 hours or upon confluency of the stamped colony, media was exchanged for mTeSR-Plus supplemented with BMP4 (50ng/mL). Colonies were allowed to differentiate in the presence of BMP4 for 48 hours before being processed for downstream analyses.

### Mesoderm Induction Media Differentiation

hiPSCs were dissociated with Accutase and resuspended in mTeSR-Plus supplemented with ROCK inhibitor, as previously described. Cells were then seeded as a monolayer at a concentration of approximately 500 cells/mm^2^. Cells were incubated at 37°C for 24 hours before being rinsed 1x with DPBS and given fresh mTeSR-Plus without ROCK inhibitor. Approximately 24 hours later, media was exchanged for Mesoderm Induction Media (MIM). Colonies were allowed to differentiate for 24-72 hours, with MIM being exchanged daily, before use in downstream analyses.

### Western Blot

Cells of each genotype were induced to form TBXT+ mesoderm with either MIM or 4uM CHIR99021 in mTeSR+ for 48 hours prior to protein isolation. Cells were washed twice with ice-cold PBS and lysed in RIPA lysis buffer (Fisher Scientific; A32965). Three replicate wells were pooled for each genotype for each differentiation condition. The protein concentration was determined using the Pierce BCA Protein Assay Kit (Life Technologies, 23227) and quantified on a SpectraMax i3 Multi-Mode Platform (Molecular Devices). following the manufacturer’s instructions. Protein (∼20-40 μg) was transferred to the membrane using the Trans-Blot Turbo Transfer System (Biorad; 1704157). The membrane was then blocked overnight at 4°C using Intercept TBS Blocking Buffer (Li-COR; 927-70001). Primary antibodies TBXT (AF2085; 1000x; Gt) and either GAPDH (ab9485; 1000x; Rb) or B-actin (ab8226, 1000x, Ms) were diluted in Intercept T20 (TBS) Antibody dilution buffer (Li-COR) at a 1:1000 ratio and incubated with the membrane overnight at 4°C. The next morning, membranes were washed in 1x TBS-T and incubated for 1 hour at RT in the dark with species-specific secondary antibodies (Rb-680; 926-68071; Gt-680 925-68074, Ms-800 926-32212 Gt-800; 926-32214) (VWR) at 1:20,000. Membranes were subsequently washed and developed using the BioRad ChemiDoc MP. Protein levels were quantified using ImageJ by first subtracting the intensity of a blank ROI from the experimental ROI, and then calculating a normalization factor by dividing the observed housekeeping intensity by the highest observed housekeeping intensity. The observed experimental signal was then divided by the lane normalization factor to generate a normalized experimental signal. Each lane from the same blot was then converted to a percentage of the highest WT normalized experimental signal on that blot.

### Immunofluorescence

hiPSCs were rinsed with PBS 1X, fixed in 4% paraformaldehyde (VWR) for 15-20 minutes, and subsequently washed 3X with PBS. The fixed cells were permeabilized and blocked in a buffer comprised of 0.3% Triton X-100 (Sigma Aldrich) and 5% normal donkey serum in PBS for one hour, and then incubated with primary antibodies diluted in antibody dilution buffer (0.3% Triton, 1% BSA in PBS) overnight (Table S5). The following day, samples were washed 3X with PBS and incubated with secondary antibodies in antibody dilution buffer at room temperature for 2 hours. Secondary antibodies used were conjugated with Alexa 405, Alexa 555, or Alexa 647 (Life Technologies) at a dilution of 1:400. Cells were imaged at 10x, 20x, or 40x magnification on an inverted AxioObserver Z1 (Zeiss) with an ORCA-Flash4.0 digital CMOS camera (Hamamatsu).

### Scratch Assay

Approximately 50k hiPSCs of each genotype were seeded into each well of a 96-well plate following the MIM induction protocol previously described. 24 hours after the addition of MIM, a scratch was made in confluent wells manually by using a p200 pipette tip. Using ZenPro software, cells were then imaged every 12 minutes across 24 hours in Brightfield at 10x magnification on an inverted AxioObserver Z1 (Zeiss) with an ORCA-Flash4.0 digital CMOS camera (Hamamatsu) at 37°C with 5% CO^2^. Images of cells were computationally segmented, and the area occupied by cells was calculated using CellProfiler at the 24hr and 48hr time points relative to initial MIM addition (0hr and 24hr of live imaging). The area occupied at the final time point was subtracted from the area at the initial time point to yield the change in area. Wells that were not confluent at the time of the scratch or wells in which the basement membrane had been removed by the scratch were omitted from the final dataset.

### Fluorescent *In situ* hybridization

PDMS stamps coated in GFR-Matrigel were applied to 3-well chamber slides with removable silicone chamber walls (Ibidi, 80381) and gastruloids were generated as described previously. Colonies were then fixed in 4% paraformaldehyde for 15-30 minutes, rinsed in PBS, and dehydrated according to the RNAscope cultured Adherent Cell Sample Preparation for RNA Multiplex Fluorescent v2 Assay (Advanced Cell Diagnostics). Slides were stored in 100% ethanol at −20°C short term until initiation of the *in situ* hybridization protocol. The *RNAscope* protocol was then performed as outlined in User Manual 323100-USM. Catalog numbers for ACDBio RNAscope probes used in this study include FGF17 (1148351-C1), RSPO3 (413701-C2), MESP1(849231), CYP26A1 (487741), and WNT5A (604921) (Table S5). Colonies were imaged using the Olympus Fluoview FV3000 Confocal Microscope or the Nikon C2 laser scanning confocal microscope equipped with a Prime 95B 25mm sCMOS camera in collaboration with the UCSF Nikon Imaging Core and Gladstone Microscopy Core.

### Cell Harvesting for Single Nuclei Multiome ATAC + RNA Sequencing

Each of the *TBXT* genotypes was differentiated, harvested, and prepared at the same time for each of the two biological replicates. Therefore, each biological replicate represents an experimental batch. For each sample within the batch, 12 micropatterns were differentiated within each well of a 24-well plate and cells from all wells on a plate were pooled, yielding a cell suspension comprising approximately 288 colonies per sample. Nuclei were isolated and ∼9,000-12,000 nuclei/sample (TBXT-KO-2 = 19,000 nuclei) were transposed and loaded onto a 10x Chromium Chip J to generate gel bead-in emulsions (GEMs) following the 10x Chromium Next GEM Single Cell Multiome ATAC and Gene Expression Kit (10x Genomics, CG000338). GEMs were processed to produce ATAC and gene expression libraries in collaboration with the Gladstone Genomics Core. Deep sequencing was performed on the NovaSeq 6000 S4 200 cycle flow cell for a read depth of >25k reads per cell (TBXT-KO-2 = 16,000 reads per cell).

### Data Processing Using CellRanger-Arc

All ATAC and GEX datasets were processed using CellRanger-Arc 2.0.0. FASTQ files were generated using the mkfastq function, and reads were aligned to the hg38 reference genome (version 2.0.0).

### Seurat Analysis

Outputs from the CellRanger-Arc count pipeline were analyzed using the Seurat package (version 4.3.0)(Butler et al., 2018; Satija et al., 2015; Stuart et al., 2019) in R (v4.2.0). Quality control filtering included the removal of outliers due to the number of features/genes (nFeature_RNA > 2500 & nFeature_RNA < 4500, nCount_RNA > 5000 & nCount_RNA < 12,500, mitochondrial percentage > 5% and mitochondrial percentage < 20%, and ribosomal percentage > 3% and ribosomal percentage < 15%). Cell cycle scores were added using the function CellCycleScoring. ScTransform v2 normalization was then performed to integrate samples based on batch with regression based on cell cycle scores and ribosomal content (vars.to.regress = c(“S.Score”, “G2M.Score”, “percent_ribo”)). Principal component analysis (PCA) was performed using the most highly variable genes, and cells were clustered based on the top 15 principal components using the functions RunUMAP, FindNeighbors, and FindClusters, and the output UMAP graphs were generated by DimPlot. The resolution parameter of 0.4 was set so that cluster boundaries largely separated the likely major cell types. Cluster annotation was performed based on the expression of known marker genes, leading to 11 broadly assigned cell types. Cells filtered out of the ArchR dataset based on doublet identification (see” ArchR Analysis” below) were removed from the Seurat dataset (final n = 23,838 cells). Differential gene expression was then performed with the function FindAllMarkers (logfc.threshold = 0.25 and min.pct = 0.1) to generate a list of top marker genes for each cluster. In pairwise comparisons of differential gene expression, positive values reflect upregulation in mutant lines, while negative values reflect upregulation in WT.

### ArchR Analysis

Indexed Fragment files generated by the CellRanger-Arc counts function served as input for the generation of sample-specific ArrowFiles (minTSS = 4 & minFrags = 1000) using the R package ArchR v1.0.2(Granja et al., 2021). ArrowFile creation also generates a genome wide TileMatrix using 500bp bins and a GeneScoreMatrix, an estimated value of gene expression based on a weighted calculation of accessibility within a gene body and surrounding locus. Each Arrow file (n=6 total) was then aggregated into a single ArchRProject for downstream analysis. Corresponding Gene Expression Matrices were imported to the project based on the filtered feature barcode matrix h5 file generated by CellRanger-arc counts and descriptive cluster labels were imported from the corresponding Seurat object based on cell barcodes. Cells filtered out of the Seurat dataset based on QC metrics previously described were also removed from the ArchR dataset. Cell doublet removal was performed in ArchR using the functions addDoubletScores and filterDoublets, leaving 23,838 cells with a median TSS of 13.278 and a median value of 12,774.5 fragments per cell. The number of cells counted as doublets and removed are found in Table S2.

After generation of the aggregated ArchR project, dimensionality reduction was performed using ArchR’s implementation of Iterative Latent Semantic Indexing (LSI) with the function addIterativeLSI based on the 500bp TileMatrix with 4 iterations, increasing resolution values (0.1, 0.2, and 0.4) each iteration. This was repeated using the Gene Expression Matrix based on 2,500 variable features, yielding “LSI-ATAC” and LSI-RNA” reduced dimensions, respectively. The two reduced dimension values were then combined using addCombinedDims to yield “LSI_Combined,” which was used as input for batch correction using Harmony with the function addHarmony (groupby = “Sample, “Batch”). Clustering was then performed using Harmony-corrected values with addClusters with a resolution of 0.4 from the R package Seurat. Finally, clusters were visualized with function plot embedding, using batch-corrected single-cell embedding values from Uniform Manifold Approximation and Projection (UMAP) using the function addUMAP. Clusters and their corresponding UMAP projection were very similar to those generated based on RNA data in Seurat, and unless otherwise stated. Cluster identities in figures are based on barcodes transferred from Seurat rather than ArchR’s LSI implementation.

After cluster annotation, pseudobulk replicates of cells within similar groups were created to facilitate peak calling. Replicates were created using addGroupCoverages and peak calling was performed using addReproduciblePeakSet using standard settings by implementing MACS2. We then used ArchR’s iterative overlap peak merging method to create a union peakset of 322,520 unique peaks.

Cluster-enriched marker peaks were identified with getMarkerFeatures, using a Wilcoxon test and normalizing for biases from TSS enrichment scores and sequencing depth, and visualized with plotMarkerHeatmap, filtering for FDR < 0.01 and abs(Log2FC) > 1.25. Motif enrichment of cluster-enriched peaks was done using addMotifAnnotations with the “CODEX” motif set. Enriched motifs per cluster were visualized by first running peakAnnoEnrichment, with FDR < 0.1 and Log2FC > 0.5. The top 7 motifs per cluster were visualized as a heatmap using plotEnrichHeatmap.

Peak-to-gene linkage analysis was performed in ArchR using the addPeak2GeneLinks command, using the batch-corrected Harmony embedding values. A total of 3,010,318 linkages were found using FDR 1e-04, corCutOff = 0.95 and a resolution of 1.

Differential accessibility within the mesoderm cluster was performed by using the command subsetArchrProject to subset the ArchR project based on the mesoderm-annotated cluster as determined from Seurat. This subsetting yielded 3,212 cells, with a median TSS of 3,085 and a median number of fragments of 12,423. Differentially expressed genes predicted pairwise across genotypes (WT vs. TBXT-KO or WT vs. TBXT-Het) were identified with getMarkerFeatures based on the GeneScoreMatrix, using a Wilcoxon Test and normalizing for biases from TSS enrichment scores and sequencing depth. GetMarkers was then run and visualized as a volcano plot using plotMarkers (FDR < 0.1 and abs(Log2FC) > 0.5). This process was repeated for the PeakMatrix to determine uniquely accessible peaks. 18 Peaks were detected between WT and TBXT-Het and 0 peaks were detected between WT and TBXT-KO. No CODEX motif enrichments were detected between genotypes (FDR < 0.1 and abs(Log2FC) > 0.5).

### CellChat

Cell signaling analysis was performed using the R package CellChat (Jin et al., 2021). The Seurat object containing all samples was subset by genotype, yielding a separate Seurat object for WT, TBXT-Het, or TBXT-KO. These 3 objects were then imported into CellChat using the function createCellChat. All ligand-receptor and signaling pathways within the CellChatDB.human were kept for analysis. Initial preprocessing to identify over-expressed ligands and receptors was performed using the functions identifyOverExpressedGenes and identifyOverExpressedInteractions with standard settings. Inference of cell communication was calculated with computeCommunProb(cellchat) and filtered by filterCommunication(cellchat, min.cells = 10). Pathway-level cell communication was calculated with computeCommunProbPathway, and aggregated networks were identified with aggregateNet, using standard settings. Network centrality scores were assigned with the function netAnalysis_computeCentrality. This workflow was run for WT, TBXT-Het, and TBXT-KO datasets independently and differential signaling analysis was then run by merging the WT, TBXT-Het, and TBXT-KO objects with mergeCellChat(). Information flow, which is defined by the sum of communication probability among all pairs of cell groups in the inferred network (i.e., the total weights in the network), was compared across genotypes using rankNet(cellchat). The distance of signaling networks between WT and TBXT-KO datasets was calculated by performing joint manifold learning and classification of communication networks based on functional similarity using computeNetSimilarityPairwise(cellchat), netEmbedding(cellchat), and netClustering (cellchat). Circle diagrams, heatmaps, and bubble plots of pathways of interest were then generated for each genotype separately using the standard settings for netVisual_aggregate(cellchat), netVisual_heatmap(cellchat), or netVisual_bubble(cellchat), respectively. Violin plots of differential gene expression were generated using plotGeneExpression(cellchat) with the standard settings.

### Gene Ontology Analysis

Gene Ontology (GO) analysis for downregulated or upregulated TBXT-dependent genes was performed with ShinyGO V0.77 (Ge et al., 2020) using GO Biological Process terms. Downregulated or upregulated TBXT-dependent gene lists from the mesoderm subcluster were assembled from differential tests between TBXT-Het vs. WT or TBXT-KO vs. WT in Seurat. Gene sets were filtered with a significance threshold set at an adjusted p-value> 0.05 and the abs(log2FC) > 0.25. Biologically relevant pathways within the first twenty hits for the TBXT-KO vs. WT comparison were visualized with bar plots, and all results are available in Table S3.

### Quantification and Statistical Analysis

Each experiment was performed with at least three biological replicates except multiomic snATAC and snRNA-seq, which was performed with two biological replicates. Multiple comparisons were used to compare multiple groups followed by unpaired *t*-tests (two-tailed) between two groups subject to a posthoc Bonferroni correction. In gene expression analysis, two replicates were used for each condition, and all gene expression was normalized to control wild-type populations followed by unpaired *t*-tests (two-tailed). Significance was specified as Adj. *P* < 0.05 unless otherwise specified in figure legends. All error bars represent the standard error of the mean (s.e.m.) unless otherwise noted in the figure legend.

## ACKNOWLEDGEMENTS

We would like to thank the Mylinh Bernardi and Horng-Ru Lin of the Gladstone Genomics Core, the Gladstone Histology and Light Microscopy Core, the Gladstone Stem Cell Core, and the Nikon Microscopy core at the University of California, San Francisco for their experimental expertise. In addition, we thank Pengfei Zhang and the Abate Lab at UCSF for assistance with PDMS stamp fabrication, as well as Nicholas Elder, David Joy, and Martin Dominguez for their experimental and computational expertise.

## AUTHOR CONTRIBUTIONS

E.A.B., and T.C.M conceived and designed the study. E.A.B, I.M.V., A.R.G.L., and B.G.B interpreted the data. E.A.B. generated the *TBXT* allelic series with input from A.R.G.L. E.A.B conducted micropattern differentiations with input from A.R.G.L. and I.M.V. E.A.B. isolated nuclei for sequencing library preparations which were processed by the Gladstone Genomics Core. E.A.B. performed immunostaining, *in situ hybridization* experiments, Western Blot, MIM differentiations, and scratch assays. E.A.B. conducted snATAC-seq, snRNA-seq analysis, including CellChat analysis. T.C.M. and B.G.B. supervised and advised. E.A.B. prepared figures and wrote the manuscript with input from all co—authors.

## DATA AVAILABILITY

snATAC-seq and snRNA-seq data have been deposited in GEO under the accession number GSE245998. Analysis scripts used to generate figure panels are freely available from the authors upon request.

## SUPPLEMENTARY FIGURES

**Figure S1:**
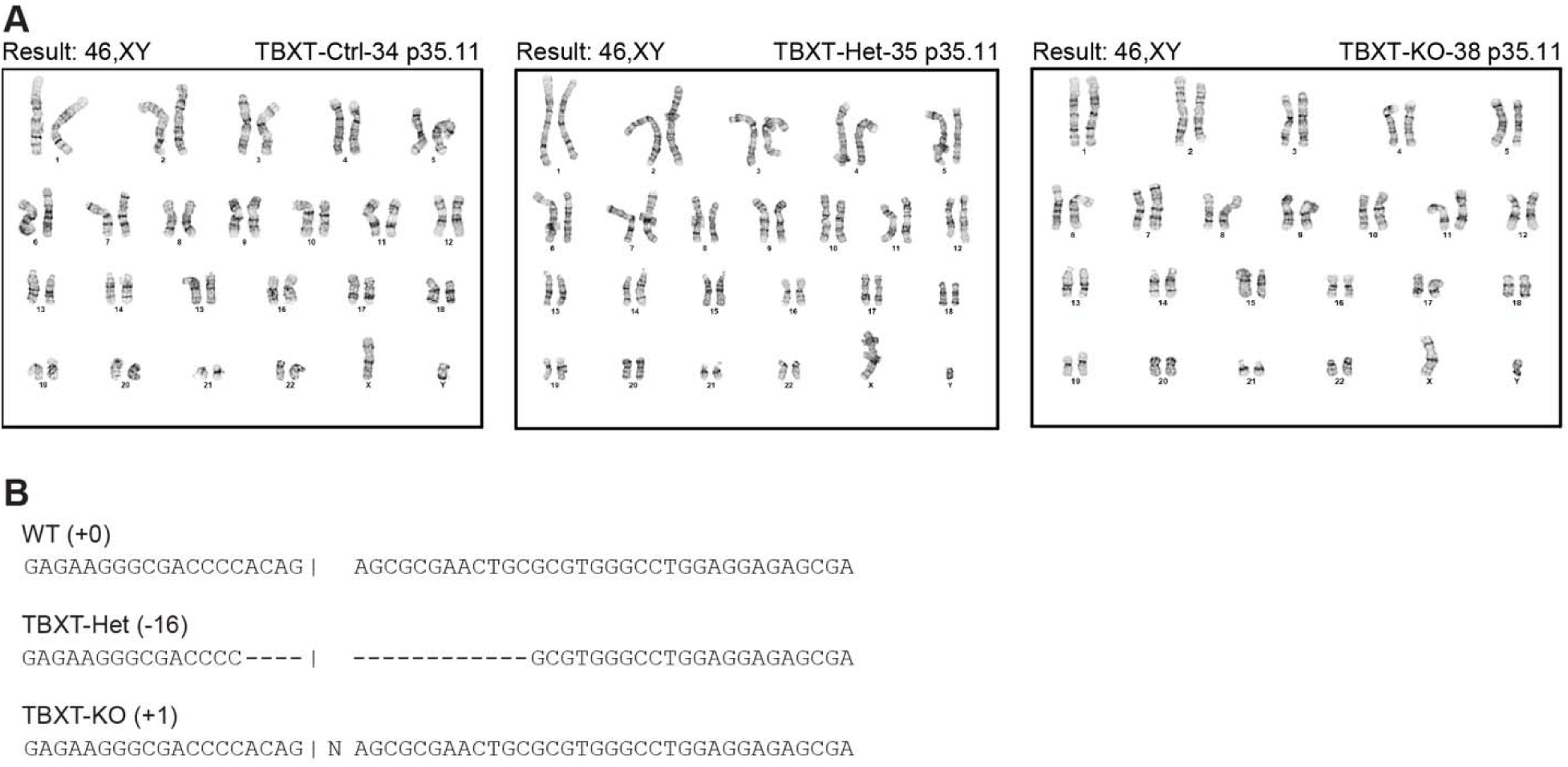
(**A**) Karyotyping results from WT (TBXT-Ctrl-34), TBXT-Het (TBXT-Het-35), and TBXT-KO (TBXT-KO-38) cell lines. (**B**) Sequences of WT, TBXT-Het, and TBXT-KO cells illustrating corresponding indels.

**Figure S2:**
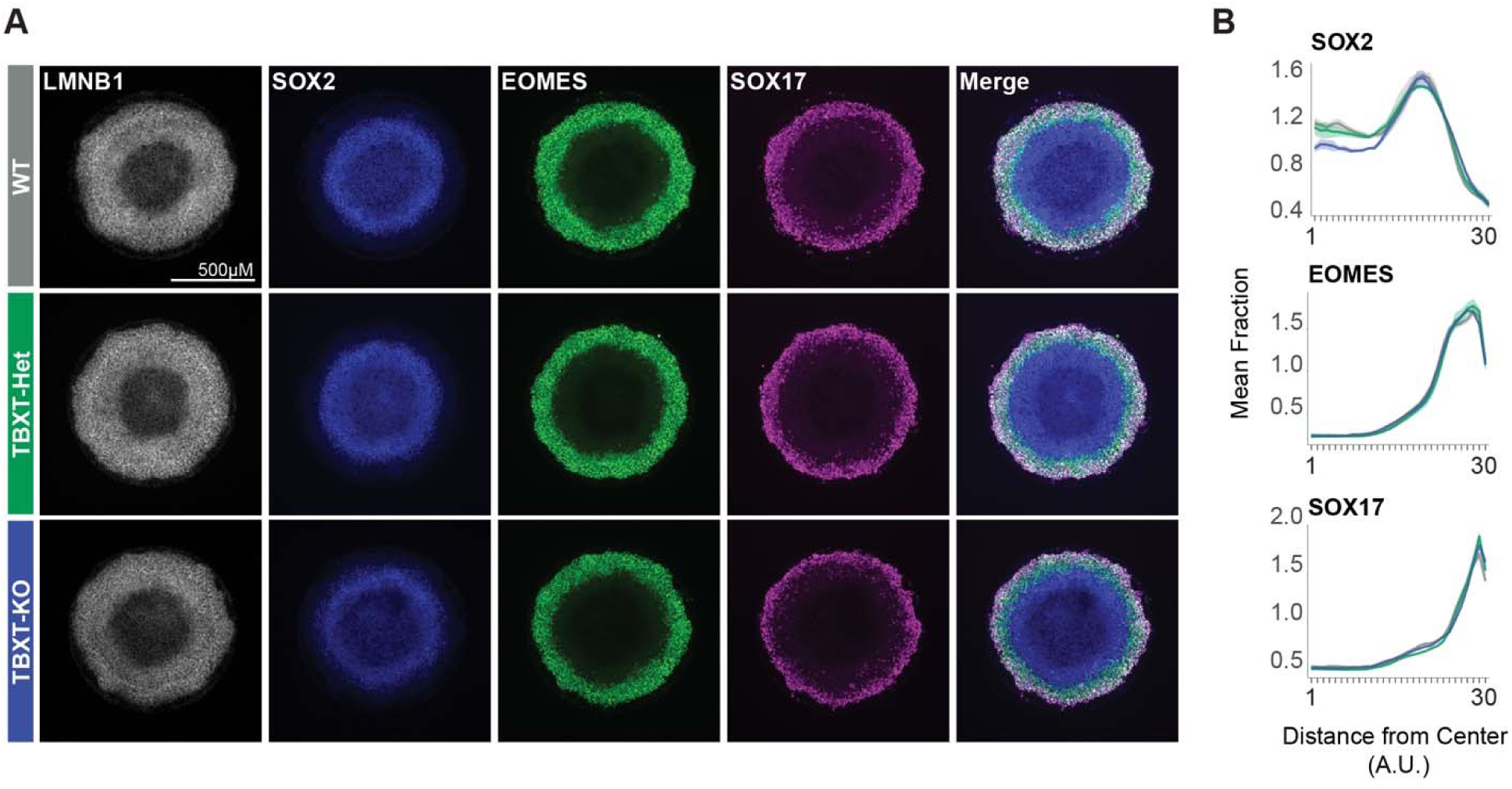
(**A**) Immunofluorescent images of 2D gastruloids for canonical gastrulation markers and (**B**) quantification of fluorescent intensity (n = 4 WT, 3 TBXT-Het, 3 TBXT-KO). Shadows around quantification data points indicate SEM.

**Figure S3:**
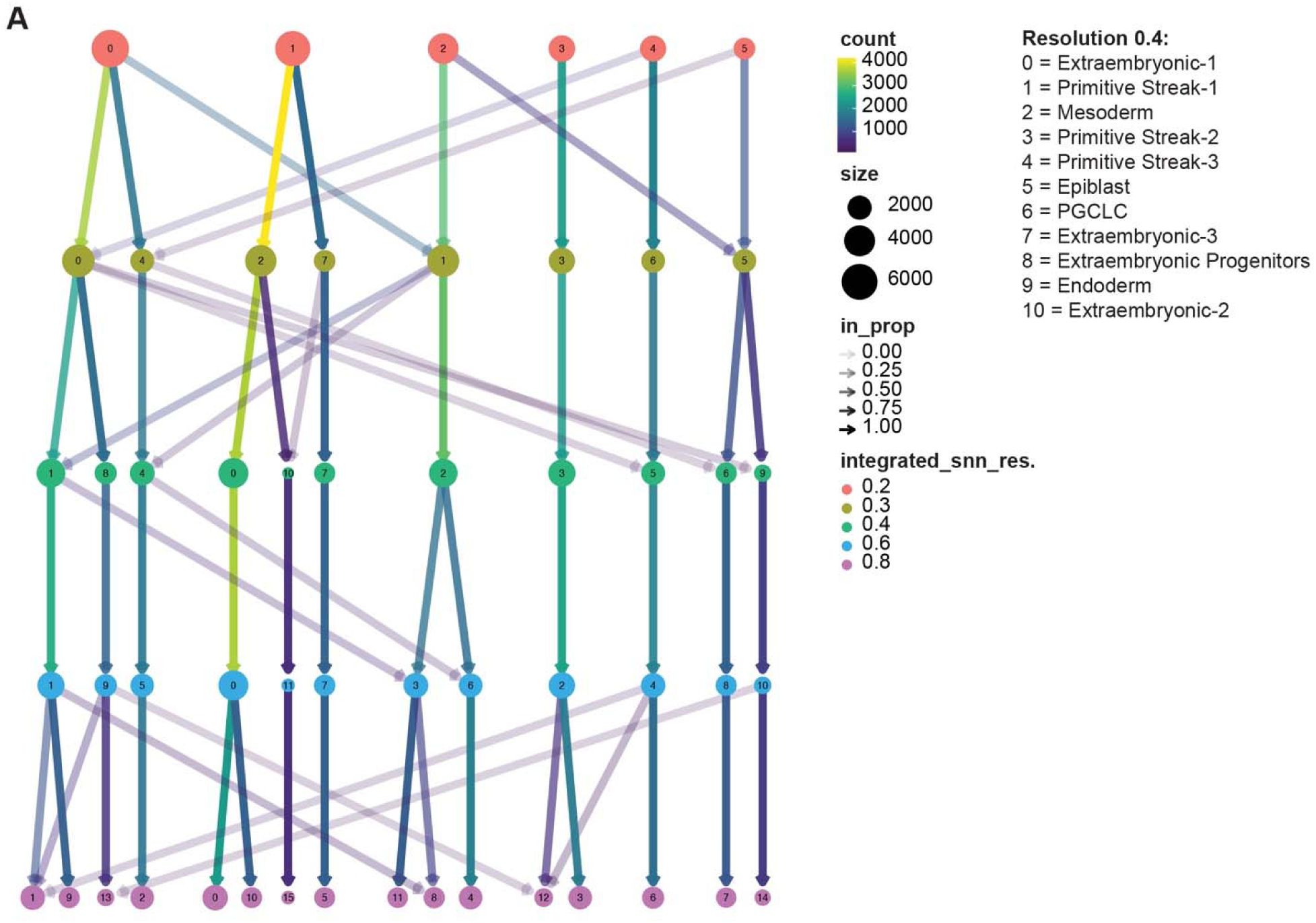
(**A**) ClusTree analysis of clusters determined at 0.2, 0.4, 0.6, and 0.8 resolution.

**Figure S4:**
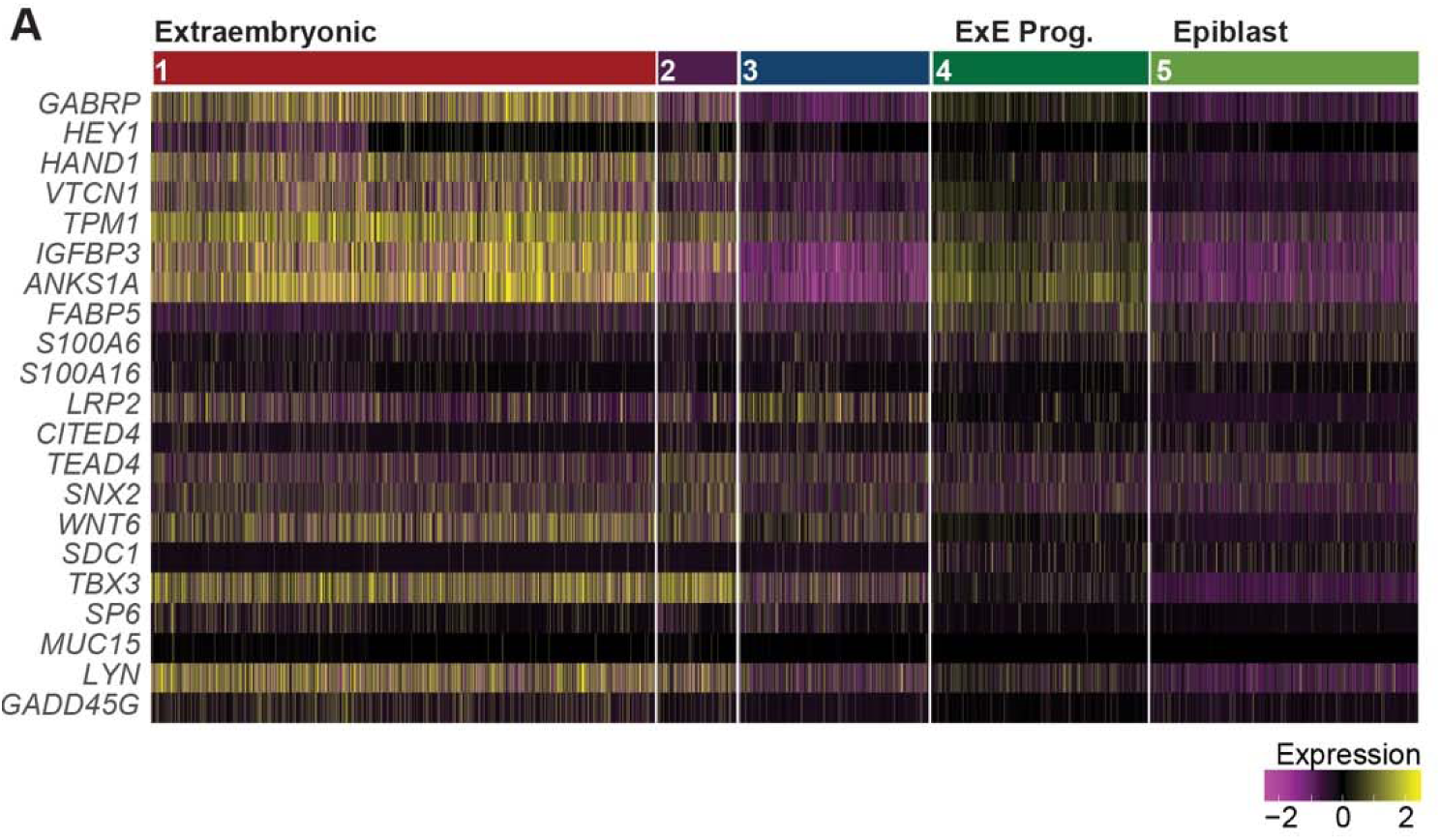
(**A**) Heatmap of extraembryonic/amnion markers for snRNA-seq data for clusters 1-5.

**Figure S5:**
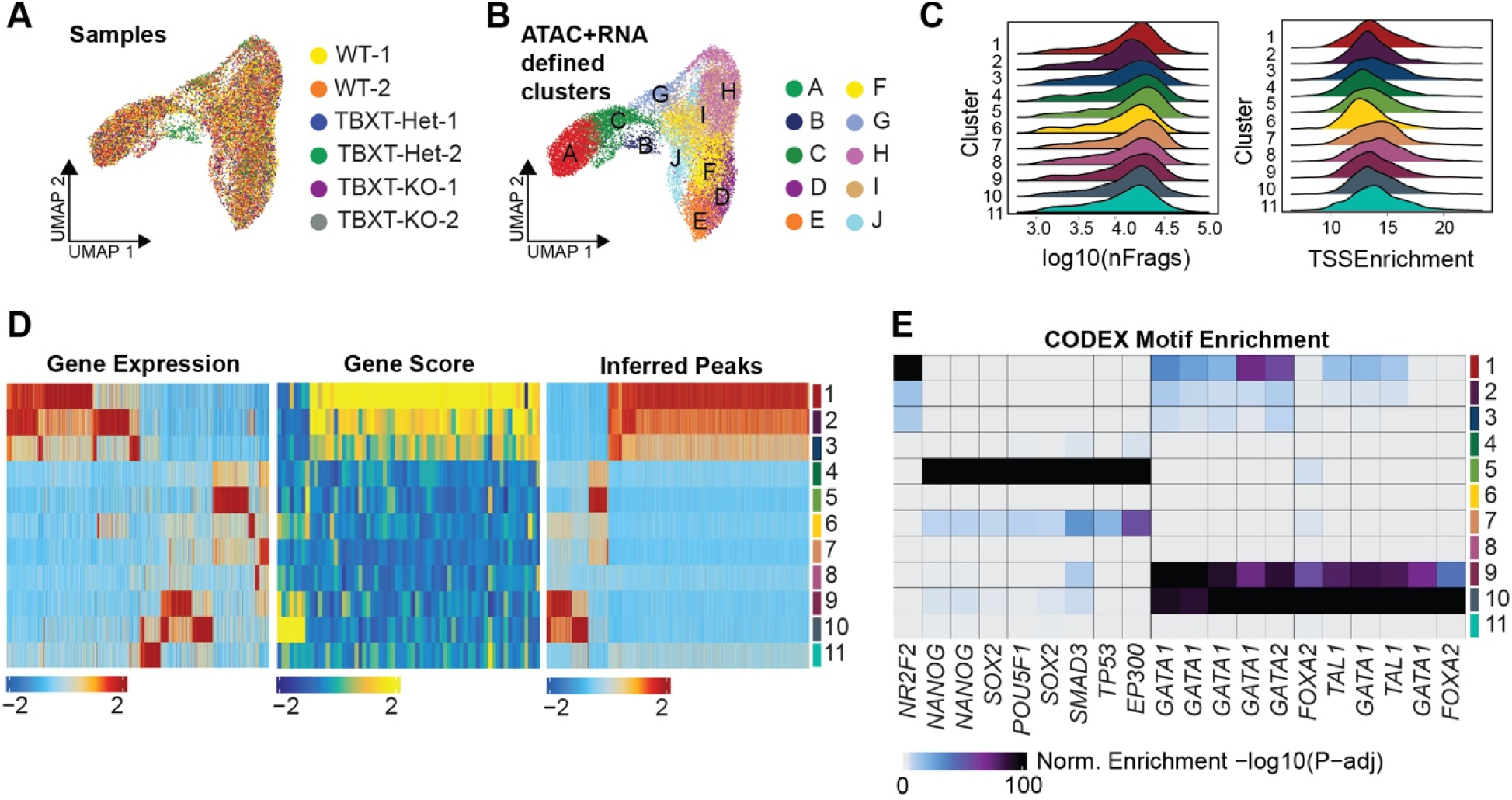
(**A**) UMAP colored by sample and (**B**) ArchR-defined clusters. ArchR clustering integrates both snRNA-seq and snATAC-seq. (**C**) Ridgeplot showing the number of fragments (log10nFrags) and TSS enrichment score across snRNA-seq defined clusters from Fig. 2. (**D**) Comparison between gene expression (GeneExpressionMatrix), predicted expression based on accessibility (GeneScoreMatrix), and inferred peaks (PeakMatrix) across snRNA-seq defined clusters from Fig. 2. (**E**) CODEX motif enrichment across snRNA-seq defined clusters from Fig. 2.

**Figure S6:**
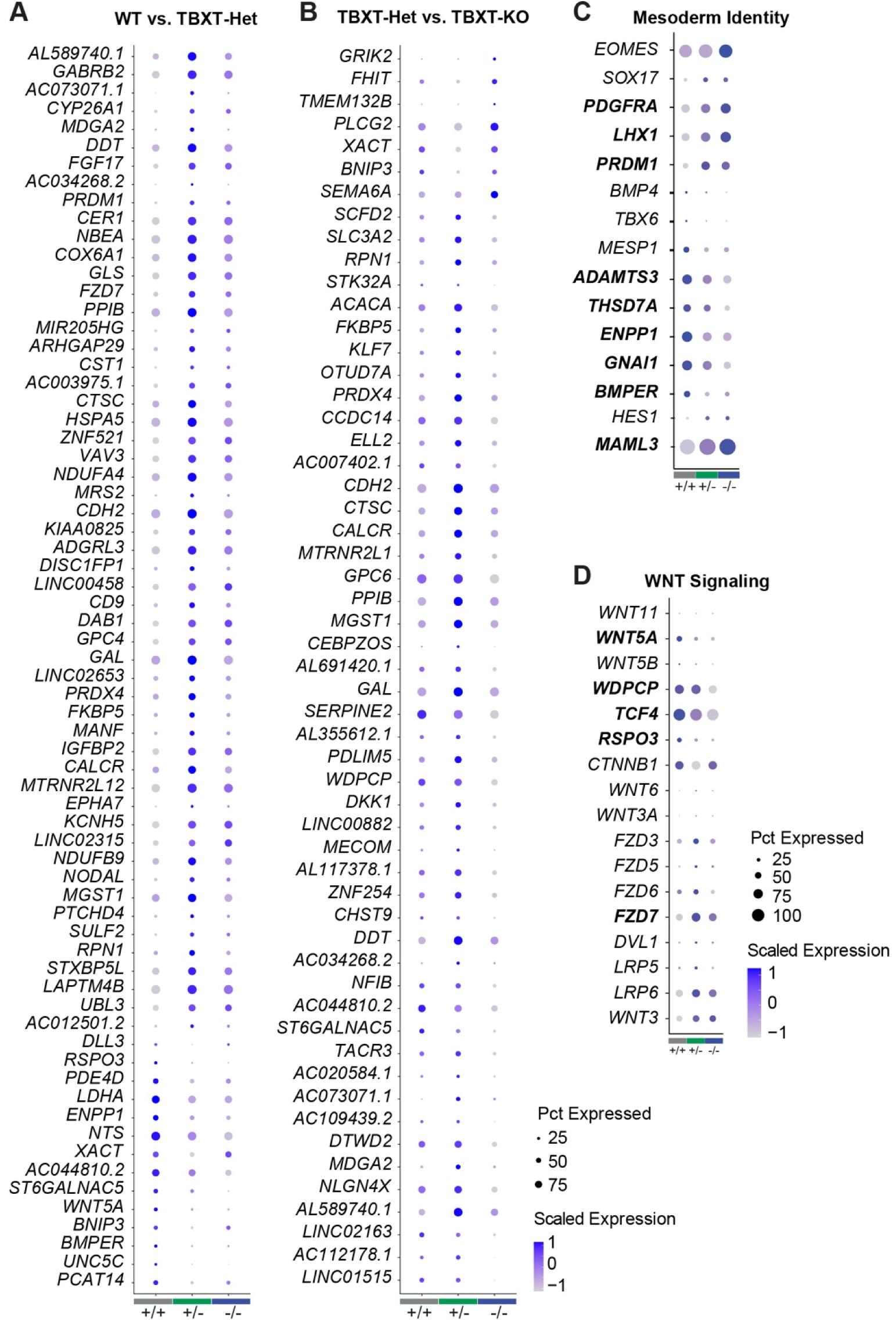
(**A**) Dot plot depicting all DE genes between WT and TBXT-Het or (**B**) TBXT-Het and TBXT-KO. (**C**) Dot plot depicting additional genes related to mesoderm identity and (**D**) WNT signaling. Bold genes in (**C-D)** indicate statistically significant DE genes (adj. p < 0.05, abs(log2FC) > 0.25).

**Figure S7:**
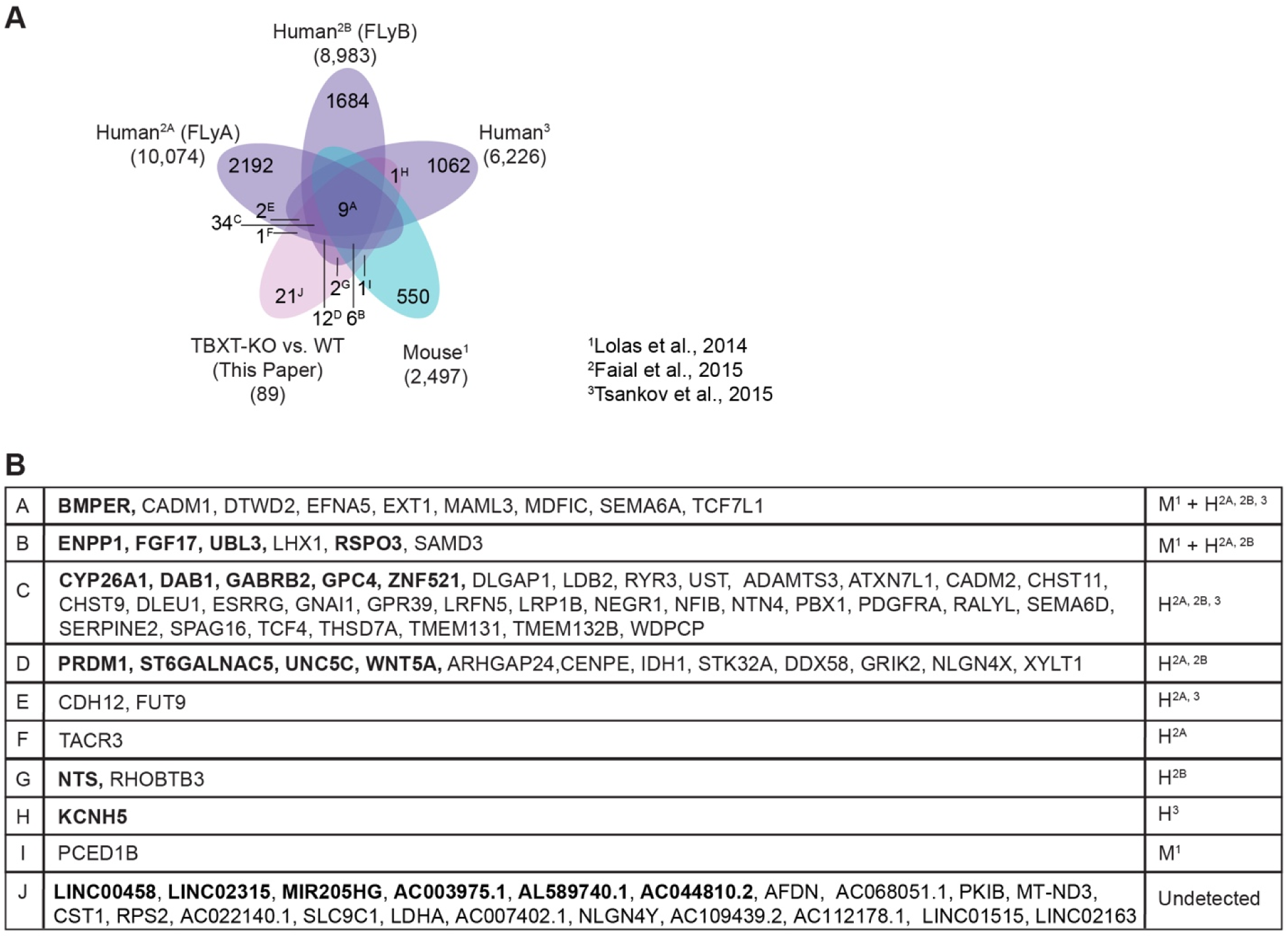
(**A-B**) Venn Diagram depicting the intersection between DE genes identified between WT and TBXT-KO and each of the 4 TBXT ChIP-seq datasets. Bold indicates significant DE genes comparing both TBXT-Het and TBXT-KO to WT (p<0.05, log2FC >= 0.25).

**Figure S8:**
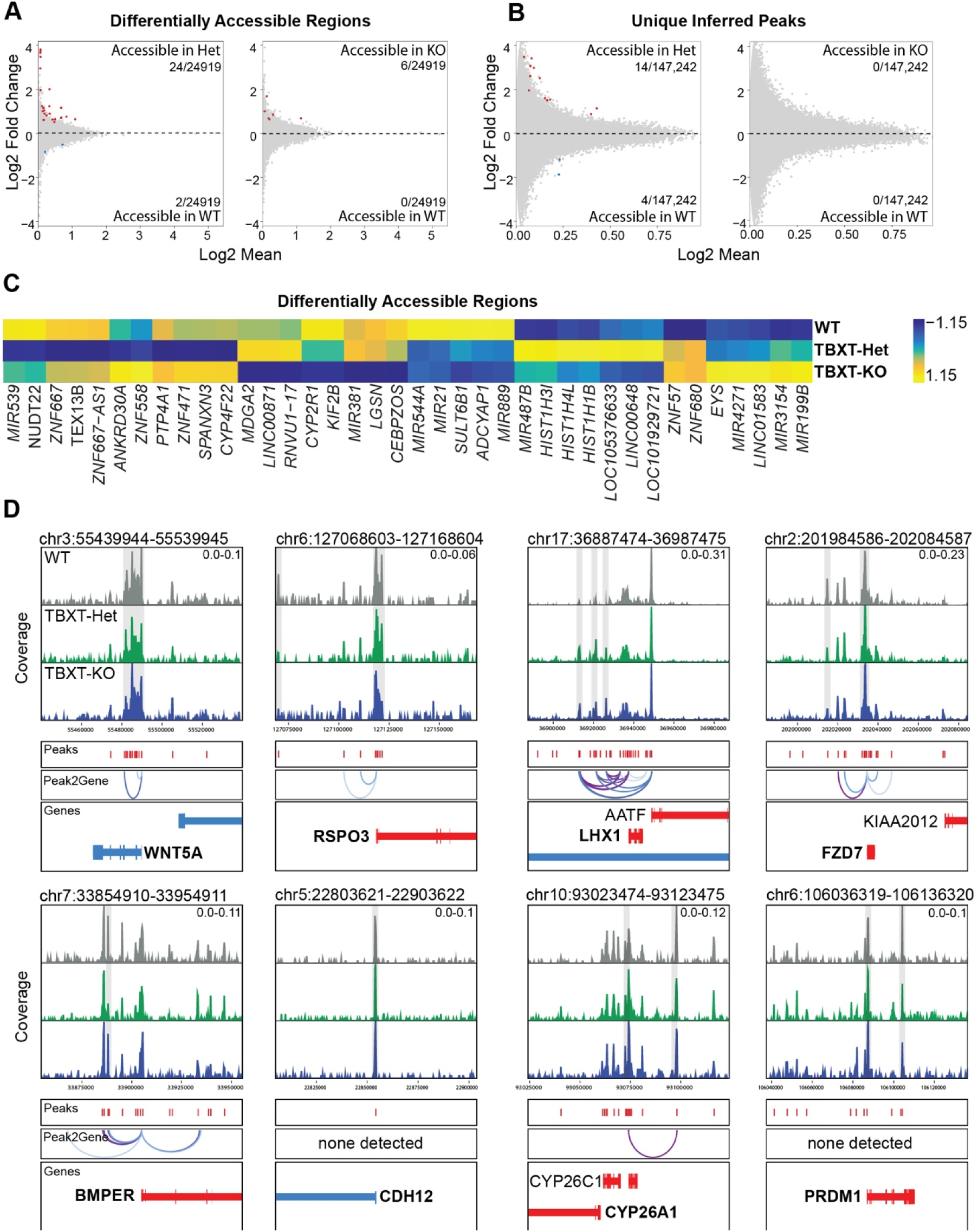
(**A**) MA plots depicting pairwise DARs and unique peaks between WT and TBXT-Het or (**B**) WT and TBXT-KO identified in the mesoderm subset (FDR < 0.1, abs(log2FC) > 0.5). (**C**) Heatmap of markerFeatures (DARs) detected across all 3 genotypes (FDR < 0.01 & abs(Log2FC) > 1.25) (**D**) Accessibility tracks comparing genotypes within the mesoderm subset centered around DE genes identified by snRNA-seq, including Peak2Gene linkage predictions of regulatory connections between distal accessible regions (peaks) and nearby genes. Gray bars indicate peaks of interest.

**Figure S9:**
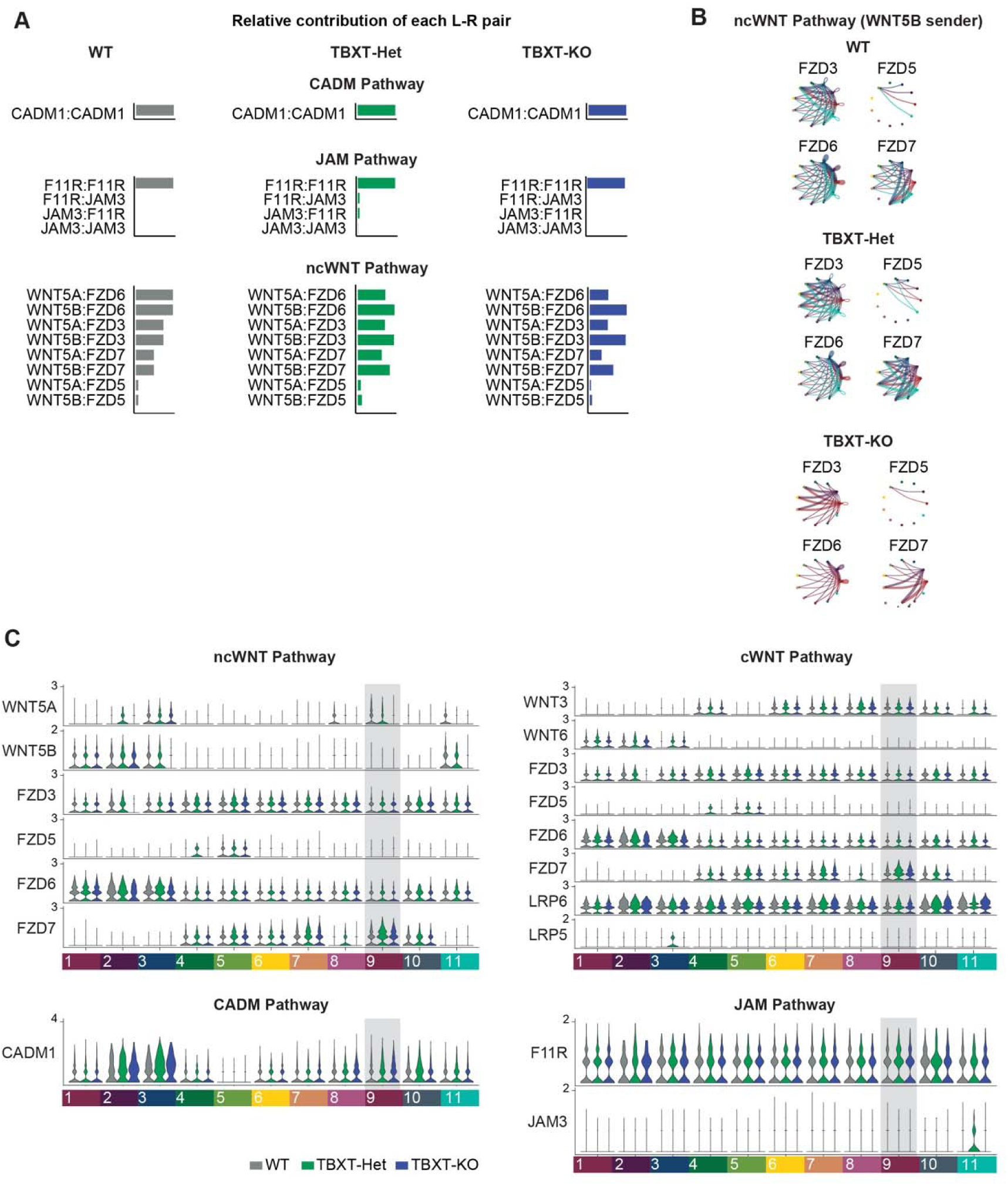
(**A**) Bar plot of the relative contribution of each ligand-receptor (L-R) pair within the CADM, JAM, and ncWNT pathways. (**B**) Circle plots visualizing predicted patterns between WNT5B and designated FZD receptors across clusters as indicated in Fig. 4. (**C**) Violin plots comparing snRNA-seq expression for key ligands or receptors between genotypes for each cluster.

**Figure S10:**
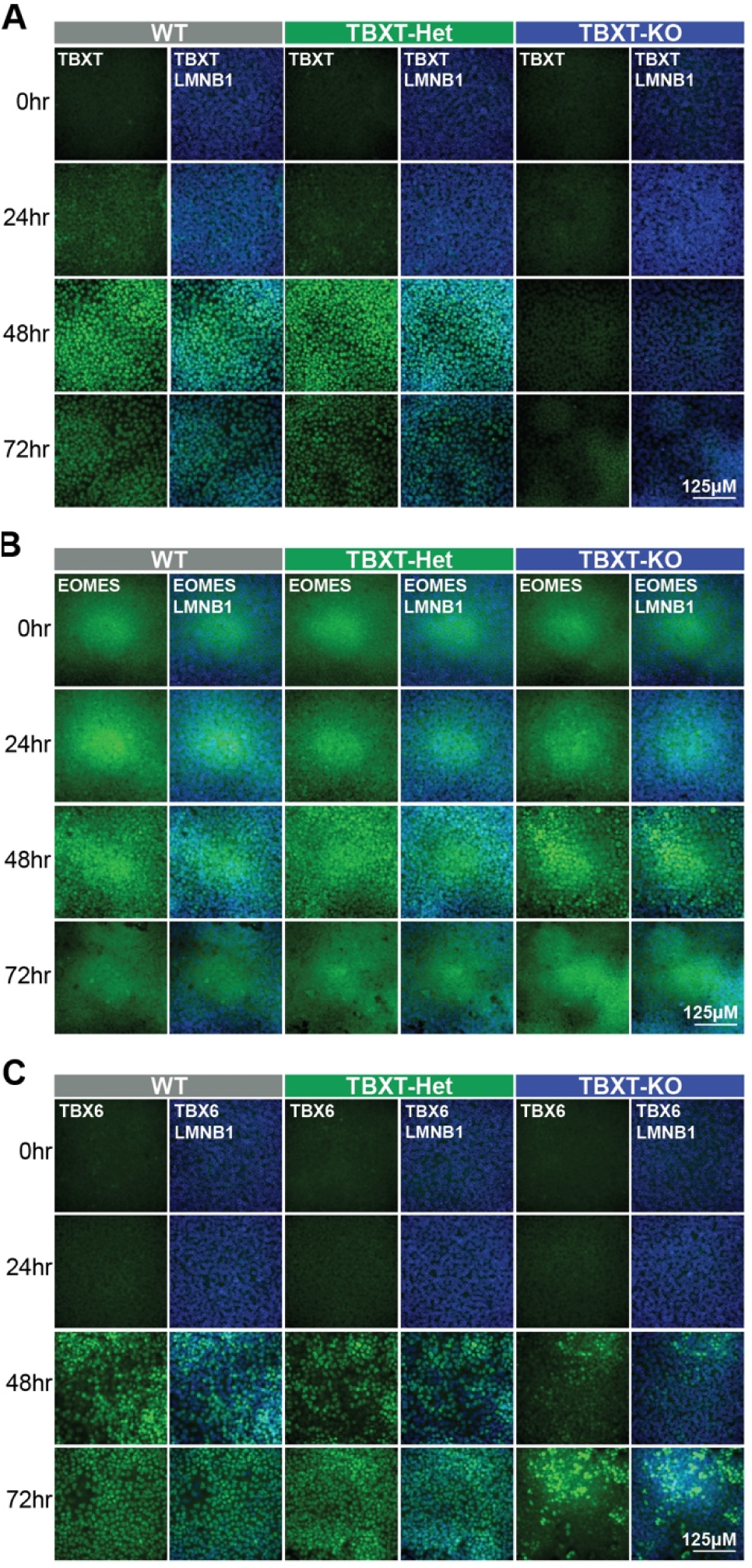
(**A-C**) Immunofluorescence for (**A**) TBXT, (**B**) EOMES, or (**C**) TBX6 at 0hr, 24hr, 48hr, and 72hr MIM exposure for each genotype. Protein of interest = green. LMNB1 = blue.

**Figure S11:**
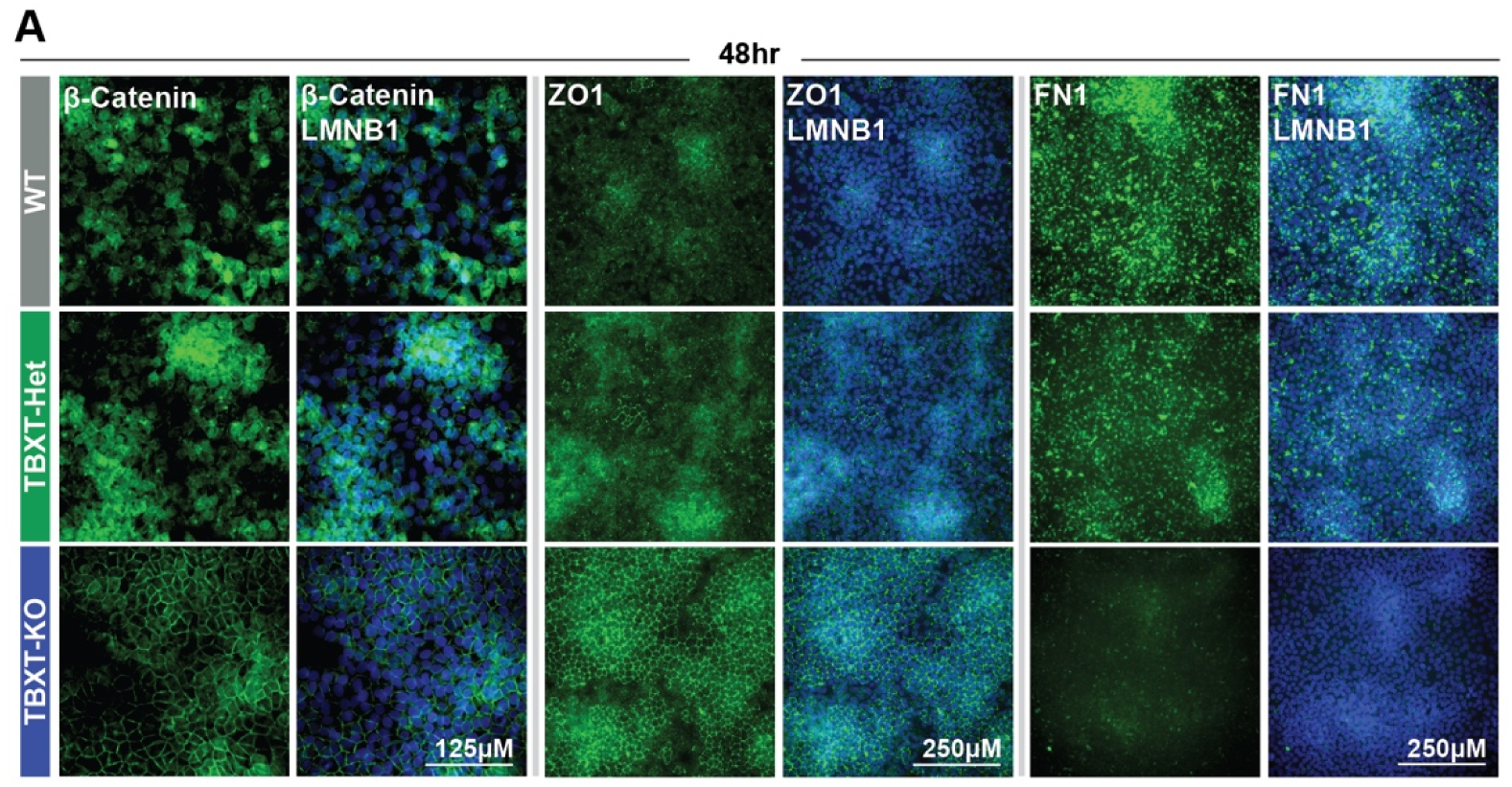
(**A**) Immunofluorescence for β –catenin, ZO1, and FN1 at 48hr MIM exposure for each genotype. Protein of interest = green. LMNB1 = blue.

## SUPPLEMENTAL TABLES

**Table S1:** Indel frequency of clonal (#34, 35, 38) or subclonal (#35-1 to 35-12) cell populations exposed to the TBXT sgRNA.

**Table S2:** Doublets filtered from each sample via ArchR analysis.

**Table S3:** ShinyGO analysis.

**Table S4:** Overview of differentially expressed genes, differentially accessible regions, and differential peaks across clusters, including unfiltered gene list for the mesoderm cluster.

**Table S5:** Antibodies used in this study.

